# Somatic Depolarization Enhances Hippocampal CA1 Dendritic Spike Propagation and Distal Input Driven Synaptic Plasticity

**DOI:** 10.1101/2021.01.12.426178

**Authors:** Tobias Bock, Steven A. Siegelbaum

**Affiliations:** Departments of Neuroscience and Pharmacology^*^, Kavli Institute for Brain Science, Zuckerman Mind Brain Behavior Institute, Columbia University Medical Center, New York, NY, 10027

## Abstract

Synaptic inputs that target distal regions of neuronal dendrites can often generate local dendritic spikes that can amplify synaptic depolarization, induce synaptic plasticity, and enhance neuronal output. However, distal dendritic spikes are subject to significant attenuation by dendritic cable properties, and often produce only a weak subthreshold depolarization of the soma. Nonetheless, such spikes have been implicated in memory storage, sensory perception and place field formation. How can such a weak somatic response produce such powerful behavioral effects? Here we use dual dendritic and somatic recordings in acute hippocampal slices to reveal that dendritic spike propagation, but not spike initiation, is strongly enhanced when the somatic resting potential is depolarized, likely as a result of increased inactivation of A-type K^+^ channels. Somatic depolarization also facilitates the induction of a form of dendritic spike driven heterosynaptic plasticity that enhances memory specificity. Thus, the effect of somatic membrane depolarization to enhance dendritic spike propagation and long-term synaptic plasticity is likely to play an important role in hippocampal-dependent spatial representations as well as learning and memory.

## Introduction

Pyramidal neurons are the principle excitatory units within the brain and process information from thousands of inputs from different brain regions to generate an action potential output. One common rule of synaptic organization is that long-range inputs from distant brain regions often target the most distal regions of a pyramidal neuron’s apical dendrites, whereas local excitatory inputs target regions closer to the soma ^1^. As a result of this dendritic organization, local and long-range inputs differentially trigger action potential output. Whereas the local inputs produce large suprathreshold somatic excitatory postsynaptic potentials (EPSPs), long-range inputs generate very small subthreshold EPSPs at the soma due to attenuation by the dendritic cable. Nonetheless distal inputs are thought to play important roles in regulating neural activity, in part, because the active properties of dendrites enable the distal inputs to trigger the firing of distal dendritic spikes ^2^. Such distally-generated dendritic spikes have been implicated in hippocampal-dependent spatial learning^3^, place field generation ^4^, and neocortical-mediated sensory perception ^5, 6^.

In the hippocampus, CA1 pyramidal neurons (PNs) receive both direct long-range connections from the entorhinal cortex that target CA1 distal dendrites and local connections from the hippocampal CA3 Schaffer collaterals (SC) that target CA1 proximal dendrites ^7^. Genetic and chemical lesion studies suggest that the distal inputs are important for temporal ^8^ and contextual memory ^9^ and play a role in fine-tuning CA1 place field firing ^10^. Although dendritic spikes help boost the influence of these distal inputs at the CA1 soma, recordings from both acute hippocampal slices and computational modeling indicate that CA1 dendrites powerfully attenuate these spikes, limiting their influence to only a small subthreshold depolarization ^11, 12^.

How can we reconcile the behavioral importance of the distal inputs with their weak somatic excitation? Even though dendritic spikes on their own produce only a small subthreshold somatic depolarization, they can interact with more proximal excitatory inputs to trigger the firing of bursts of somatic action potentials termed complex spikes ^11, 13^. Active dendritic spikes also provide an important source of Ca^2+^ entry for generating long-term synaptic plasticity at distal synapses ^13, 14^. Finally, dendritic spikes have been proposed to serve as instructive signals by mediating a heterosynaptic form of plasticity termed input-timing-dependent plasticity (ITDP), in which dendritic spikes triggered by distal inputs produce a long-term enhancement of CA1 excitation by the SC inputs ^15, 16^, thereby enhancing the specificity of contextual memory ^9^.

In addition to being regulated by coincident proximal synaptic stimulation, several lines of evidence suggest that distal synaptic inputs may exert a surprisingly large influence on somatic output under in vivo conditions. Thus, CA1 neurons show relatively normal rates of place cell firing when proximal SC inputs are lesioned ^17^ or genetically silenced ^18^, which has been attributed to a high efficacy of the remaining influence of the distal inputs. What might be responsible for the apparent greater efficacy of distal synaptic inputs in triggering somatic output under in vivo conditions compared to ex vivo slices?

Here we examine the potential influence of somatic resting potential on the generation and propagation of distal dendritic spikes since the soma often experiences significant subthreshold depolarizations in vivo as a result of excitatory input. The importance of somatic voltage for in vivo neural activity has been previously established by the finding that small depolarizations from the resting potential promote the firing of complex spikes^19^ and lead to the de novo appearance of CA1 place field firing in formerly silent neurons^20^. Larger depolarizations promote the firing of long-lasting dendritic plateau potentials that generate de novo place fields^4^, presumably by triggering synaptic plasticity.

However, the mechanism by which somatic depolarization may amplify distal synaptic inputs remains incompletely understood. A recent study found that in rat hippocampal slices, somatic depolarization can amplify the response at the CA1 neuron soma to a subthreshold burst of distal EPSPs as a result of the activation of a perisomatic persistent voltage-dependent Na^+^ conductance^21^. A previous study reported that somatic depolarization also enhances the propagation of suprathreshold dendritic spikes in CA1 pyramidal neurons in isolated rat hippocampal slices^22^. However, the mechanism by which weak somatic depolarization alters dendritic spike propagation and dendritic-spike-dependent forms of plasticity remains unknown. Here we report that somatic depolarization in mouse CA1 pyramidal neurons powerfully enhances the propagation of distal dendritic spikes, but not subthreshold EPSPs, by enhancing the resting inactivation of dendritic A-type voltage-gated K^+^ channels. Furthermore, somatic depolarization dramatically enhances the efficacy of induction of ITDP, an effect that is due, at least in part, to the enhancement of dendritic spike propagation, providing a potential mechanism for regulating the specificity and strength of hippocampal-dependent declarative memory.

## Results

### Dendritic spike propagation efficacy is modulated by the somatic membrane potential

To explore the regulation of dendritic spike propagation in hippocampal CA1 PNs, we obtained dual whole cell current clamp recordings from the soma and distal apical dendrite of these cells, ∼250-300 μm from the soma (at the border of stratum radiatum and stratum lacunosum moleculare), in acute hippocampal slices from adult mice (Fig. 1a). We triggered a distal dendritic spike by injecting a 350 pA, 100 ms depolarizing current step through the dendritic patch pipette and recorded the resultant voltage change at the soma (Fig. 1b). In the dendrite, the resultant spike had a fast rising component, typically associated with voltage-gated Na^+^ channels ^23, 24^, followed by a slower plateau potential, typically generated by voltage-gated Ca^2+^ channels ^23, 25, 26, 27^. When the somatic membrane potential was held initially at -70 mV, near the typical CA1 resting potential, the dendritic spike (32.4 ± 5.3 mV in amplitude) produced only a very small somatic depolarization (0.65 ± 0.08 mV, n = 5, Fig. 1b-d), for a dendritic attenuation factor (dendrite/soma spike amplitude) of 49.8 ± 5.7, consistent with previous findings ^11, 12^.

**Figure 1:**
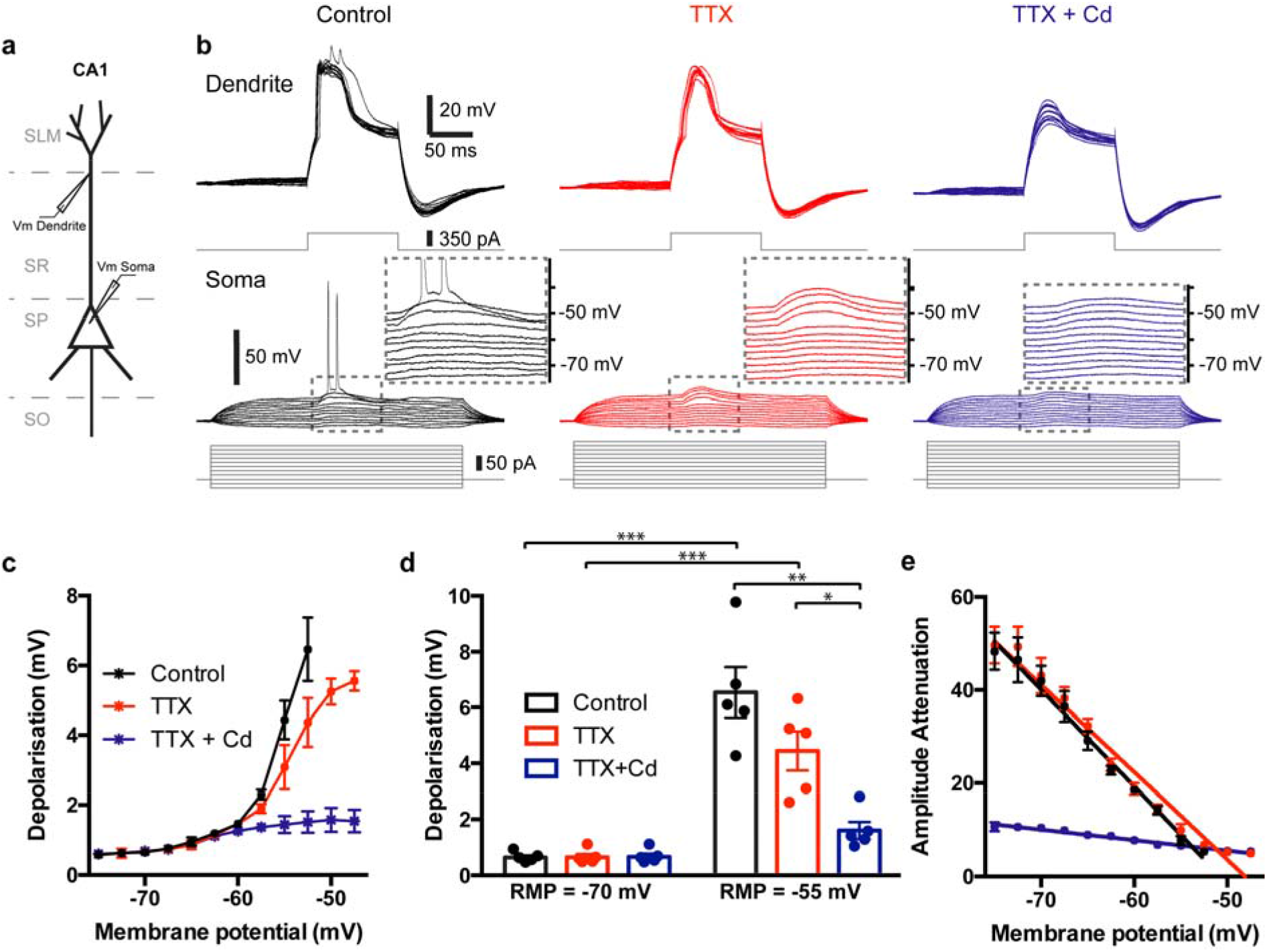
Somatic resting membrane potential determines the efficacy of dendritic spike propagation to the soma. **a** Schematic of the experimental setup. **b** Dendritic (top) and somatic (bottom) recording of a dendritic spike, elicited by current injection into the dendrite at varying somatic membrane potentials before (black) and after adding 500 nM TTX (red) followed by 200 µM Cd (blue) to the bath solution; inserts show an expanded view of the traces within the dashed box. **c** Peak depolarization of the somatic membrane potential in response to a dendritic spike at varying somatic baseline potentials before (black) and after adding TTX (red) and Cd (blue) to the bath. **d** Comparison of peak somatic membrane depolarization at -70 mV and -55 mV baseline somatic membrane potential before (black) and after adding TTX (red) and Cd (blue) to the bath. **e** Attenuation factor of the dendritic calcium spike, as it propagates to the soma, plotted against the somatic baseline potential before (black) and after adding TTX (red) and cadmium (blue) to the bath.

However, depolarization of the somatic resting potential by constant current injection dramatically enhanced the efficacy of dendritic spike propagation by nearly an order of magnitude. Thus, with the soma held initially at -55 mV, a distal dendritic spike produced a 6.46 ± 0.90 mV somatic depolarization, decreasing the attenuation factor to 7.79 ± 1.19 (p < 0.005, n = 5; Fig. 1b-d). Upon further depolarization of the soma, a dendritic spike typically elicited a burst of somatic action potentials (Fig. 1b). Changes in somatic membrane potential had no impact on dendritic spike threshold, amplitude or duration at the distal dendritic recording site (Supplementary Fig. 1). Somatic depolarization also caused no significant change in the passive properties of the cell in either compartment (Supplementary Fig. 1). The signal attenuation factor between the distal dendritic recording site and the soma showed an S-shaped dependence on membrane potential, decreasing continuously to an asymptotic value as the somatic membrane potential was depolarized (p < 0.0001 with Friedman test, n = 5, Fig. 1e). We term this phenomenon depolarization-enhancement-of-dendritic-spike-propagation (DEDSP).

To determine the ionic nature of the dendritic spikes and the mechanism by which somatic depolarization enhanced their propagation, we examined the effects of antagonists of voltage-gated Na^+^ and Ca^2+^ channels. Inhibition of voltage-gated Na^+^ channels with tetrodotoxin (TTX, 500 nM) blocked the fast phase of the dendritic spike but left the slower plateau potential (indicative of Ca^2+^ spikes ^23, 24, 25, 26, 27^) intact. Somatic depolarization enhanced the propagation to the soma of these Ca^2+^ spikes in a manner similar to that seen under control conditions. In TTX the somatic amplitude of the dendritic spike was 0.66 ± 0.13 mV at a somatic resting potential of -70 mV compared to 4.36 ± 0.69 mV at a resting potential of -55 mV (p < 0.05, n = 5, Fig.1b-d), indicating that DEDSP does not require voltage-gated Na^+^ channels. Combined application of the voltage-gated Ca^2+^ channel blocker Cd^2+^ (200 μM) and TTX fully eliminated the dendritic spike, even in the distal dendrite (0.69 ± 0.11 mV at -70 mV versus 1.51 ± 0.31 mV at -55 mV, p = 0.07, Fig.1b-d), confirming that the plateau potentials were Ca^2+^ spikes. Importantly, with dendritic spikes blocked, somatic depolarization had only a minimal effect on propagation of the passive dendritic depolarization.

### Modulation of dendritic spike propagation efficacy during somatic membrane depolarization is dependent on A-type K^+^ channels

Previous studies have implicated voltage-gated A-type Kv4.2/4.3 K^+^ channels in regulating dendritic excitability. These channels are present in the apical dendrites of CA1 pyramidal neurons where they regulate the back-propagation of dendritic spikes ^28^, curtail the duration of dendritic plateau potentials, and confine their spread throughout different dendritic branches ^29^. Moreover, these channels rapidly inactivate over a voltage range that matches the effect of somatic depolarization we observed ^28, 30, 31^, providing a potential mechanism for how depolarization may enhance dendritic spike propagation.

To examine directly the role of A-type K channels in DEDSP, we compared the effects of membrane depolarization on dendritic spike propagation before and after bath application of the A-type K-channel blocker, 4-aminopyridine (4-AP, 5 mM). Application of 4-AP greatly enhanced spike propagation at a somatic holding potential of -70 mV, and largely occluded the enhancement in spike propagation with membrane depolarization (Fig. 2). Although some residual effect of depolarization on propagation efficacy remained in the presence of 4-AP (p < 0.005 with Friedman test, Fig. 2b,d), non-linear fits to the data showed that 4-AP significantly diminished the effect of voltage (p < 0.01 with extra-sum-of-squares-F Test, Fig. 2b,d). In the presence of 4-AP, the slope of a linear regression of the dendritic attenuation curve (Figure 2d) was reduced by 93.8%, from -1.94 to -0.12 (p < 0.0001). Furthermore, while somatic depolarization from -70 mV to -55 mV caused a large and highly significant increase in the amplitude of the dendritic spike at the soma (0.97 ± 0.2 mV compared to 4.94 ± 0.78 mV; p < 0.005, n = 5; Fig. 2c), depolarization caused only small, non-significant increase in the somatic response in the presence of 4-AP (4.42 ± 0.43 mV vs. 5.79 ± 0.55 mV; p = 0.063, n = 5, Fig. 2c). Finally, 4-AP produced a similar enhancement of dendritic Ca^2+^ spike propagation in the presence of TTX, confirming that voltage-gated Na^+^ channels are not required (Fig. 2b-d).

**Figure 2:**
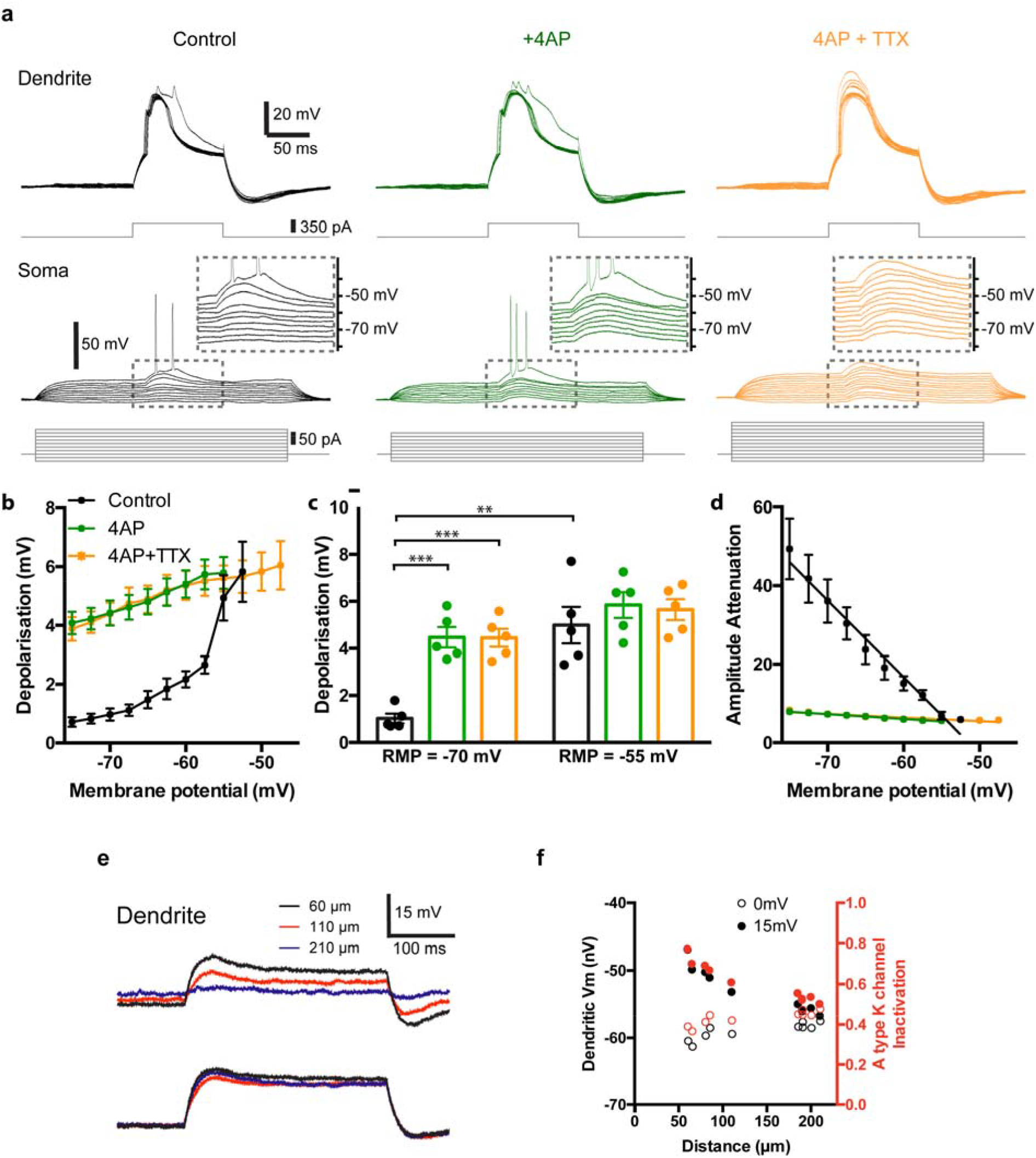
The voltage dependence of the propagation efficacy of dendritic spikes is mediated by 4-AP-and phrixotoxin-sensitive K^+^ channels. **a** Dendritic (top) and somatic (bottom) recording of a dendritic spike, elicited by current injection into the dendrite at varying somatic membrane potentials before (black) and after adding 5 mM 4-AP (green) and 500 nM TTX (orange) to the bath solution, inserts show a zoom of the traces of somatic membrane potential within the squares. **b** Peak depolarization of the somatic membrane potential during the dendritic spike at varying somatic baseline potentials before (black) and after adding 4-AP (green) and TTX (orange) to the bath. **c** Comparison of peak somatic membrane depolarization at -70 mV and -55 mV baseline somatic membrane potential before (black) and after adding 4-AP (green) and TTX (orange) to the bath. **d** Attenuation factor of the dendritic calcium spike, as it propagates to the soma, plotted against the somatic baseline potential before (black) and after adding 4-AP (green) and TTX (orange) to the bath. **e** Voltage traces at the soma and different dendritic locations in response to a 15-mV somatic depolarization (black: 60 µm distance from soma, red: 110 µm distance, blue: 200 µm distance). **f** Depolarization (black) with estimated A-type K^+^ channel availability (1 – steady-state inactivation) shown in red at various dendritic locations when the soma is depolarized by 15 mV (closed circles) or when the soma is not depolarized (open circles).

As 4-AP is also known to block Kv1 delayed-rectifier channels ^32^, we examined the effect of the specific Kv4.2/3 blocker phrixotoxin ^33, 34^. Under control conditions, the somatic depolarization in response to a dendritic spike increased significantly from 1.26 ± 0.36 mV at a somatic membrane potential of -70 mV to 4.56 ± 0.26 mV when the somatic membrane was depolarized to -55 mV (p < 0.005, n = 5, Supplementary Fig. 2f-h). Similar to the action of 4-AP, 500 nM phrixotoxin caused a significant increase in the amplitude of the somatic response to a dendritic spike at a somatic resting potential of -70 mV (4.28 ± 0.37 mV; p<0.005 versus control in absence of phrixotoxin; n=5). Moreover, the enhancement in dendritic spike propagation normally seen upon depolarization of the membrane to -55 mV was largely abolished by the toxin (5.12 ± 0.41 mV at – 55 mV, p = 0.094, n = 5, Supplementary Fig. 2h), with the slope of the best linear fit to the somatic response–voltage relation reduced by 91.6%, from 1.67 to 0.14 (p < 0.0001 with linear regression test; Supplementary Fig. 2h).

As neither 4-AP nor phrixotoxin altered dendritic passive membrane properties, dendritic spike threshold, amplitude or duration (Supplementary Fig. 2), we surmised that somatic depolarization might enhance dendritic spike propagation by enhancing the resting inactivation of A-type K^+^ channels. To test this idea, we measured the effects of somatic depolarization on the proximal dendritic membrane voltage of CA1 pyramidal neurons to determine whether the extent of depolarization was sufficient to alter channel inactivation. We performed multiple dual patch clamp recordings from the soma and from the proximal dendrites at various distances from the soma. Consistent with previous observations on spatial attenuation of voltage changes ^35, 36^, we saw a strong depolarizing effect of a 15 mV somatic depolarization on the proximal dendrites, but little effect in the distal dendrites (Fig. 2e,f). Using the inactivation versus voltage plot for A-type K^+^ channels from Hoffman et al. (1997) and correcting for liquid junction potential (−6.2 mV), we found that the measured effect of somatic depolarization should markedly increase steady-state inactivation of the A-type channels in the proximal dendrites within 100 - 150 μm of the soma (Fig. 2f). Thus, in the proximal dendrite (60 µm), resting inactivation of A-type K^+^ channels was predicted to increase from 38% when the soma was at -70 mV to 78% when the soma was depolarized to -55 mV. In contrast, in the distal dendrite (>200 μm from the soma) we observed only a small increase in resting inactivation from 45% to 50% (Fig. 2f). From this data, we conclude that somatic depolarization results in a marked increase in resting inactivation of A-type K^+^ channels in the proximal apical dendrite that allows for more efficient propagation of dendritic spikes during somatic depolarization, while distal dendritic locations (the site of dendritic spike initiation) are barely affected by alterations in somatic voltage.

### Somatic depolarization enhances propagation of synaptically-evoked dendritic spikes but not subthreshold PSPs

Next we investigated whether somatic depolarization can also enhance the propagation of dendritic spikes generated in a more physiological manner through synaptic stimulation of the perforant path (PP) inputs to the distal CA1 dendrites. We placed an extracellular stimulating electrode in the stratum lacunosum-moleculare (SLM) of CA1 (at least 200 μm from the dendritic patch location), the distal dendritic site of the PP inputs, and applied either a single stimulating pulse to elicit a subthreshold PSP or a strong, brief high-frequency burst of stimuli to elicit a dendritic spike (4 or 5 stimuli at 100 Hz, see Fig. 3a).

**Figure 3:**
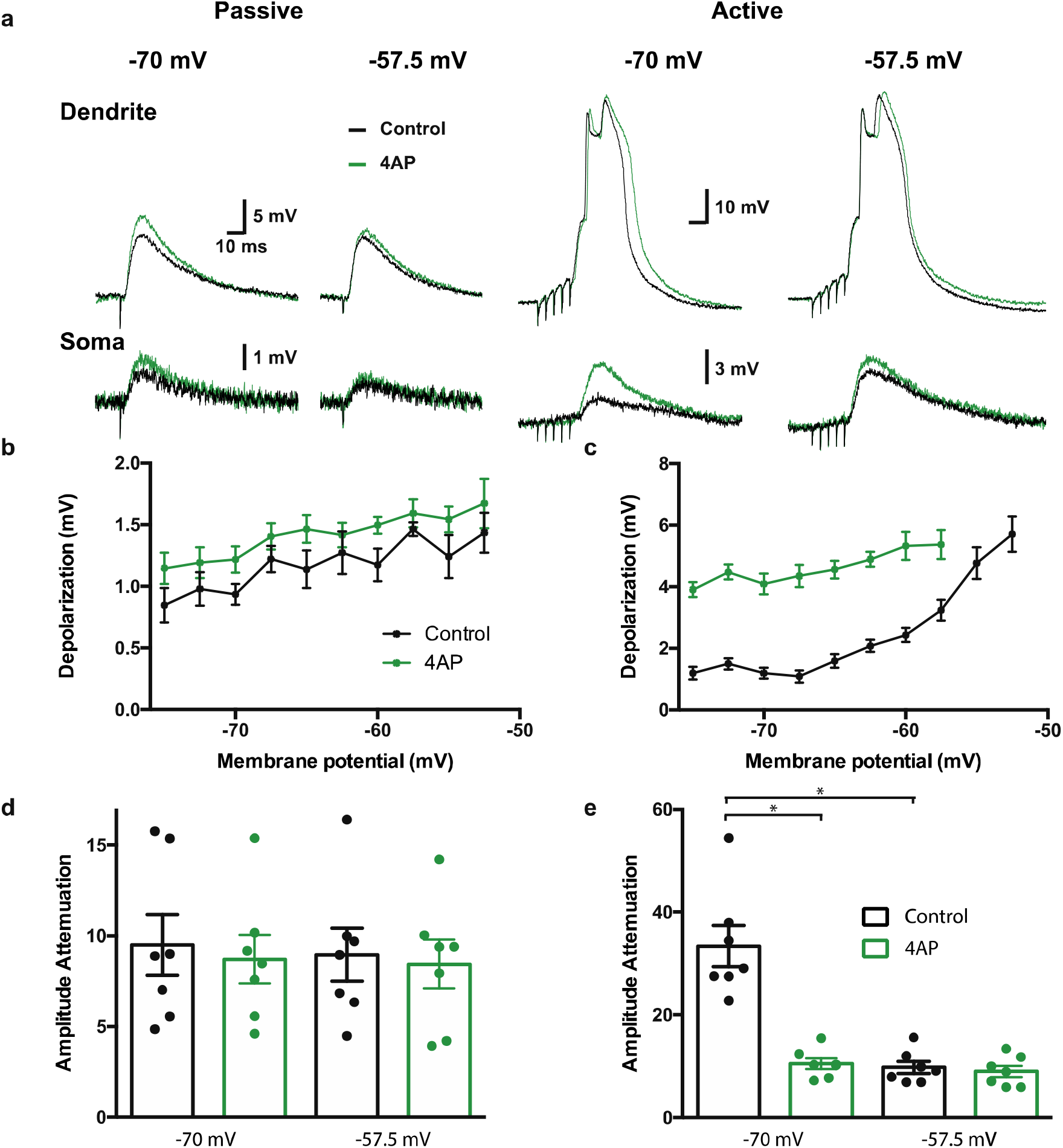
A-type K-channels affect active and passive signals differently. **a** Example traces of dendritic and somatic responses to single PP stimulation, which evokes a passively propagated PSP (left) and a train of 5 pulses of PP stimulation at 200 Hz, which evokes an actively propagating dendritic spike (right) when the soma is held at wither -70 or -57.5 mV before (black) and after (green) application of 5 mM 4-AP. **b** Peak depolarization of the somatic membrane potential during a passively propagating PSP at varying somatic baseline potentials before (black) and after adding 4-AP (green). **c** Peak depolarization of the somatic membrane potential during an actively propagating dendritic spike at varying somatic baseline potentials before (black) and after adding 4-AP (green). **d** the attenuation factor of passively propagating dendritic signals if the soma is held at -70 mV (left) or at -57.5 mV (right) before (black) or after (green) application of 4-AP. **e** Same as **d** but for actively propagating signals.

Propagation of the subthreshold PSP evoked by a single perforant path stimulus showed little dependence on somatic depolarization. Thus the PSP amplitude at the soma was attenuated by a factor of 9.49 ± 1.67 at -70 mV and 8.96 ± 1.45 at -57.5 mV (p = 0.94, n = 7, Fig. 3a-d). In contrast, the propagation of a dendritic spike evoked by a brief train of synaptic stimuli (5 stimuli at 200 Hz) was greatly enhanced by somatic depolarization in a manner similar to that observed when we evoked a dendritic spike through direct current injection (see Fig. 1). Thus, the synaptically-evoked spike amplitude attenuation at the soma was 33.4 ± 4.0 at -70 mV versus 9.8 ± 1.2 at -57.5 mV, p = 0.015, n = 7, Fig. 3a,c,e). Moreover, with sufficient somatic depolarization, the synaptically evoked dendritic spike also led to burst firing of somatic APs.

The application of 4-AP (5 mM) caused a small but significant increase in the subthreshold PSP amplitude in both the dendrite and the soma (in both cases p = 0.03, n = 7, Fig. 3a and Supplementary Fig. 3) but did not alter dendritic propagation of the subthreshold PSP, at either a somatic voltage of -70 mV or -57.5 mV (Fig. 3b, d; dendritic attenuation was 8.71 ± 1.34 at -70 mV and 8.44 ± 1.34 at -57.5 mV, p = 0.57, n = 7, Fig. 3a,b,d). In contrast, the propagation of the synaptically–evoked dendritic spike was greatly enhanced by bath application of 4-AP at more negative but not depolarized somatic membrane potentials. In 4-AP, spike attenuation at -70 mV (10.53 ± 1.05; n = 7) was significantly less than that in control conditions (33.4 ± 4.0; p = 0.0156, Fig. 3e). In contrast, at -57.5 mV attenuation in 4-AP (9.02 ± 1.10) was similar to that in the absence of the drug (9.78 ± 1.19, p = 0.109, n = 7, Fig. 3e). 4-AP had no significant impact on the spike amplitude in the distal dendrite (p = 0.08, n = 7, Supplementary Fig. 3) nor did it alter PSP or dendritic spike duration, indicating that A-type potassium channels are more important in regulating spike propagation along the apical dendrites in stratum radiatum than spike generation in distal dendrites in stratum lacunosum-moleculare. This is consistent with previous pharmacological results ^29^ and the pattern of expression of Kv4.2 ^37^.

### Depolarization enhances dendritic spike firing and somatic bursting during paired activation of distal and proximal synaptic inputs

Paired activation of distal and proximal synaptic inputs can synergistically interact to generate firing of action potential bursts and to induce Hebbian^13^ and non-Hebbian forms of synaptic plasticity, including ITDP ^9, 15, 16^. ITDP is selectively induced when paired stimuli are delivered at 1 Hz for 90 s and when the distal input precedes the proximal input by precisely 20 ms (−20 ms pairing interval). During the standard 90 s induction protocol from a somatic potential of -70 mV, paired stimulation initially fails to elicit a dendritic spike. However, as the train of paired stimuli continues, the probability of spike firing increases, reflecting some unknown form of short-term intrinsic or synaptic plasticity. At a resting potential of -70 mV, the dendritic spikes fail to elicit somatic action potentials^9^. This contrasts to the in vivo situation where dendritic plateau potentials trigger a burst of somatic action potential output^19^. As the CA1 PN somatic resting potential in vivo is depolarized by 10-15 mV relative to that in ex vivo slices, we examined the influence of somatic depolarization on dendritic spike firing and propagation in response to paired distal and proximal stimuli at the -20 ms ITDP pairing interval.

We obtained dual apical dendritic and somatic whole cell patch clamp recordings and either blocked A-type K-channels or manipulated the somatic membrane potential during the 90 s period of 1 Hz paired stimulation (Fig. 4a,b). As described previously, with the membrane at -70 mV a 1 Hz train of paired stimuli of distal and proximal inputs elicited dendritic spikes that propagated poorly to the soma (Fig. 4b)^9^. Application of 4-AP at a resting potential of -70 mV or depolarization of the membrane to -55 mV enhanced the propagation of dendritic spikes elicited by paired synaptic stimuli (Fig. 4e). In addition, either depolarization or 4-AP increased the probability that a pair of stimuli would elicit a dendritic spike at various times during the train (Fig 4c). More specifically, the rate of increase of dendritic spike firing throughout the time course of ITDP induction, measured as the slope of the linear regression (0.36 when the soma was held at -70 mV; n = 5) was enhanced, by both 4-AP (to 0.52, p = 0.038 compared to -70 mV with multiple comparison test; n = 5) and somatic depolarization to -55 (to 0.67; p = 0.021, compared to -70 mV with multiple comparison test; n = 5; Fig. 4c). With the membrane at -70 mV the number of dendritic spikes elicited by the last 10 paired stimuli of the train increased from 3.8 ± 0.4 mV in the absence of 4-AP to 6.4 ± 0.5 at -70 mV in the presence of 4-AP (p = 0.079). Depolarization to -55 mV in the absence of 4-AP caused an even larger increase in the number of dendritic spikes, to 8.2 ± 0.8 (p = 0.016 relative to -70 mV without 4-AP, Fig. 4d).

**Figure 4:**
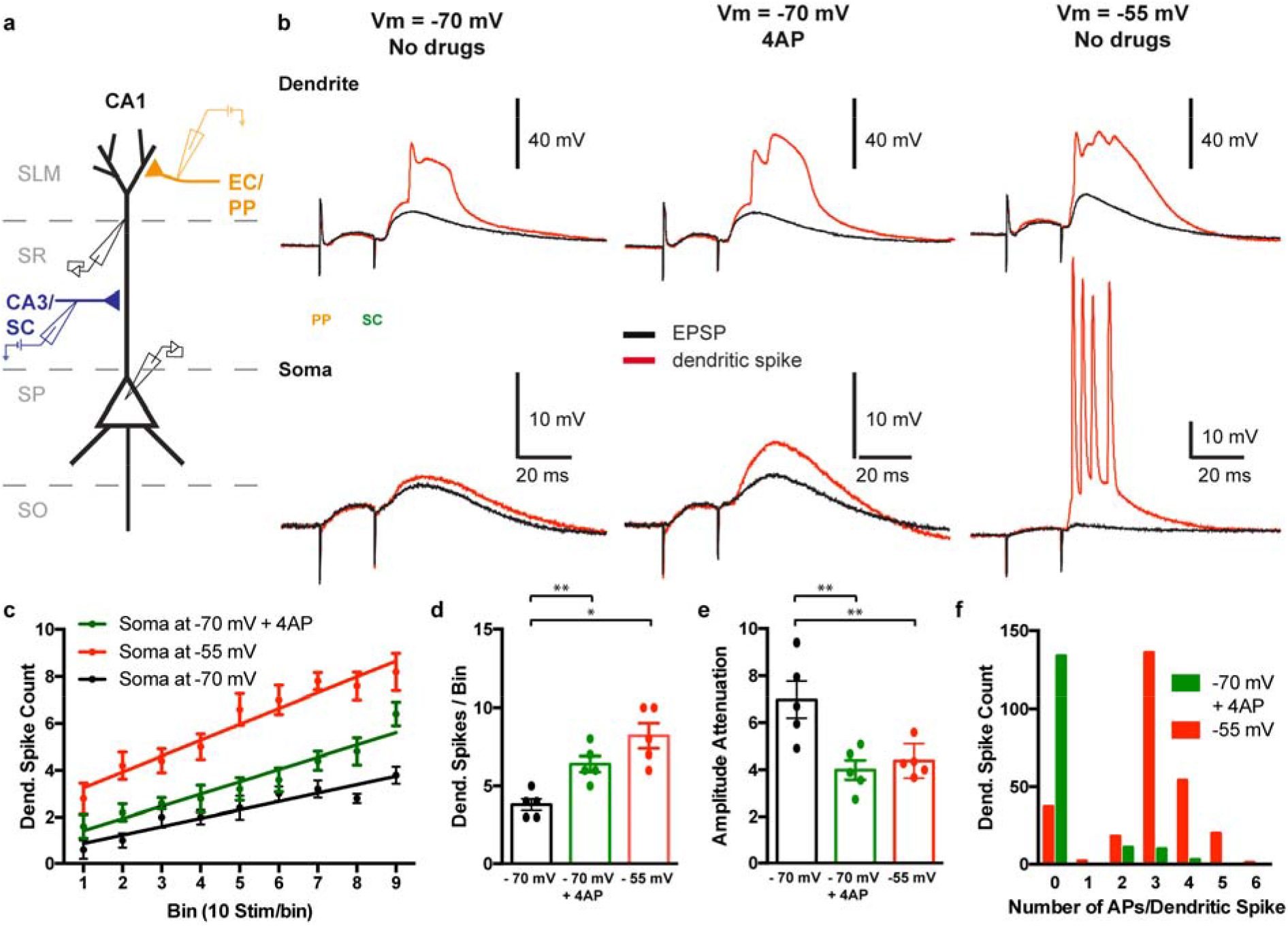
The impact of dendritic spikes at the soma during paired PP-SC stimulation. **a** Schematic of the experimental setup, showing the somatic and dendritic patch pipettes (black) as well the PP (orange) and SC (blue) stimulating electrodes. **b** Example traces of the somatic (bottom) and dendritic (top) responses to ITDP induction stimuli for all three tested conditions (left: Soma was held at -70 mV, middle: 4-AP was present in the bath during induction, right: Soma was held at -55 mV during induction stimulation), black traces show examples of subthreshold PSPs, red traces show examples in which the SC stimulus elicited a dendritic spike. **c** Average number of dendritic spikes in 10 s bins during the induction period. The 90 paired stimuli were separated into 9 bins of ten stimuli each and the number of dendritic spikes in each bin was counted (black: Soma was held at -70 mV, green: 4-AP (5 mM) was present in the bath during induction, red: Soma was held at -55 mV during induction stimulation). Straight lines show linear regression fits. **d** Dendritic spike count during the last 10 stimuli of the induction (black: Soma was held at -70 mV, red: Soma was held at -55 mV during induction stimulation, green: 4-AP was present in the bath during induction). **e** Amplitude of the somatic depolarization when a dendritic spike was elicited by the paired PP-SC stimulation (black: Soma was held at -70 mV, green: 4-AP was present in the bath during induction, red: Soma was held at -55 mV during induction stimulation). **f** Distribution of the number of somatic APs caused by dendritic spikes during paired PP-SC stimulation when 4-AP was present in the bath (green) or when the soma was depolarized to -55 mV during the induction period (red).

Of interest, when somatic APs were blocked by local application of TTX to the soma while the soma was depolarized to -55 mV, the number of dendritic spikes still increased significantly during the 90 s of paired stimulation (Supplementary Fig. 5c) and increased relative to dendritic spike firing at a somatic potential of -70 mV. Thus, the short term plastic process that enhances dendritic spike firing during the train of paired stimuli does not require somatic action potential firing but rather may be triggered by dendritic spiking alone. Moreover, as neither somatic depolarization nor 4-AP alters the local threshold in the dendrite for firing a dendritic spike in response to a single current injection (Supplementary Figures 1 and 2), the effect of 4-AP or somatic depolarization to enhance the probability of dendritic spike firing during the train of paired stimuli (relative to spike firing with the normal resting potential of -70 mV in the absence of 4-AP) likely reflects their action to enhance dendritic spike propagation.

**Figure 5:**
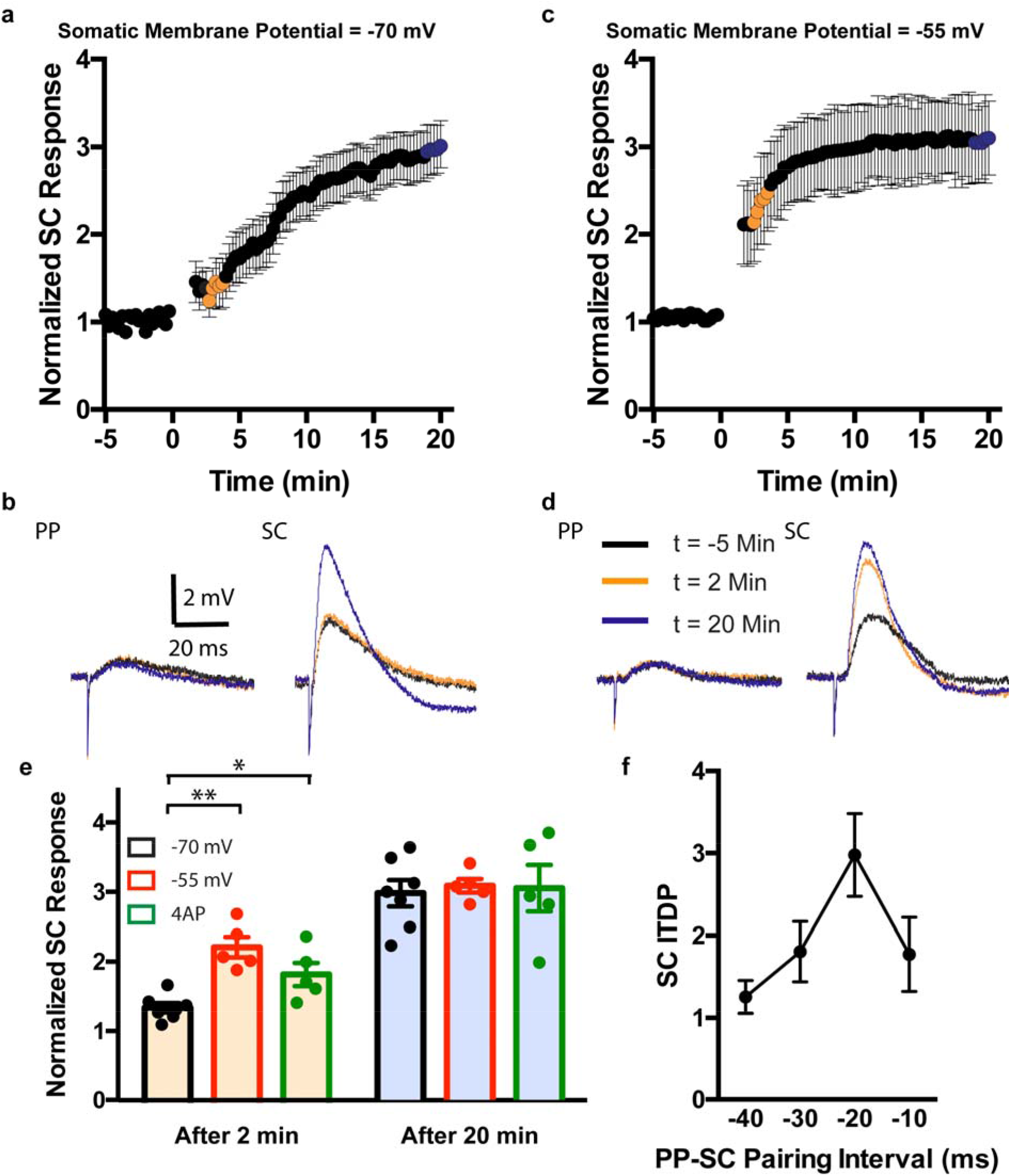
Manipulating the somatic membrane potential affects onset of ITDP expression. **a** Somatic PSP response to SC stimulation after ITDP induction, normalized to the response before induction (orange and blue sections represent the time points that were used for averages in later panels). **b** Example traces of PSPs in response to perforant path (PP, left) and SC stimulation before (black), 2 minutes (orange) and 20 minutes (blue) after ITDP induction. **c, d** Same as **a** and **b** respectively except that in this case, the soma was depolarized to -55 mV during the induction stimuli. **e** Comparison between the normalized response to SC stimulation 2 minutes (orange filled bars) and 20 minutes (blue filled bars) after ITDP induction at a membrane potential of -70 mV (black) or -55 mV (red) or if 4-AP was bath applied during ITDP induction (green). **f** Normalized response to SC stimulation 20 minutes after ITDP induction at a membrane potential of -55 mV at varying PP-SC intervals during the induction.

Although the probability of dendritic spike firing increased progressively during the 90 s of paired stimulation even with the resting membrane at -70 mV under control conditions (Fig. 4c), the dendritic spikes were ineffective in eliciting a somatic action potential. Of the 104 spikes dendritic spikes elicited by synaptic pairing, only two triggered a somatic action potential (1.9%, n = 5 cells). Moreover, in these two instances, the soma fired only one or two action potentials. Application of 4-AP or somatic depolarization greatly increased the probability that a dendritic spike would elicit one or more somatic action potentials. Thus, in the presence of 4-AP, 24 out of 158 dendritic spikes elicited a somatic AP (15.2%; p = 0.0004 with chi square test relative to control, n = 5 cells).

Moreover, when a dendritic spike was successful in activating the soma, it now triggered a burst of 2 or more spikes, with an average of 2.67 ± 0.14 somatic APs per dendritic spike. Depolarization to -55 mV was even more effective in triggering a burst of somatic actions potentials, with 231 out of 268 dendritic spikes elicited by paired stimulation evoking one or more somatic APs (86.2%, p<0.0001 with chi square test relative to control and 4-AP, n = 5 cells), with an average of 2.87 ± 0.08 APs per burst (p < 0.0001 with Mann-Whitney test relative to 4-AP, Fig. 4f).

At least two factors may contribute to the effect of somatic depolarization to enhance the firing of somatic APs in response to a dendritic spike elicited by paired stimulation. The first is the effect of somatic depolarization to enhance dendritic spike propagation. The second is the effect of somatic depolarization to bring the membrane closer to threshold. We believe that both factors are important as somatic spikes were never triggered by paired stimuli that failed to trigger a dendritic spike, even with the soma held at -55 mV. Additionally, 4-AP caused a significant increase in the amplitude of the somatic response to a dendritic spike (5.93 ± 0.53 mV to 9.39 ± 1.24 mV, p < 0.05 with Mann-Whitney test, n = 5, Fig. 4e), while not significantly changing somatic AP threshold (see Supplementary Fig. 2e), indicating that the enhanced propagation of dendritic spikes itself leads to increased firing of somatic APs.

### Manipulations that enhance dendritic spiking enhance the induction of ITDP

Our finding that somatic depolarization enhanced dendritic spike firing during paired distal and proximal synaptic stimulation suggests that somatic resting potential may also regulate the induction or expression of ITDP. When the membrane was depolarized to -55 mV, the ITDP protocol induced a similar final level of potentiation to 312.4 ± 9.5 % (n=5) of its initial level to that seen with the membrane held at -70 mV (297.2 ± 19.6 %, n=7; p = 0.76 relative to –55 mV). However, membrane depolarization greatly accelerated the time course of onset of potentiation. With the soma held at - 70 mV, the ITDP induction protocol resulted in a slowly developing enhancement in the PSP amplitude over a period of 20 min, as previously described^9, 15, 16^, with the PSP increasing to only 138.5% ± 8.1% (n=7) of its initial value two min after the end of the ITDP induction protocol. However, with the membrane held at -55 mV, the PSP increased to 225% ± 11.2 % (n=5) of its initial value at this time (p = 0.0025 relative to -70 mV; Fig. 5a-e). Application of 4-AP (with the membrane at -70 mV) also sped the onset of ITDP, with the PSP increased to 184.6 ± 14.5% (p = 0.014 relative to -70 mV with no 4-AP; n = 5) after 2 min (Fig. 5e), with no enhancement in ITDP magnitude after 20 min (PSP increased to 308.5 ± 32.7 %, p = 0.88, n = 5). Importantly, membrane depolarization did not alter the tuning of ITDP induction to the -20 mV pairing interval (p = 0.0019 with Kruskal-Wallis test, n = 5/time interval, Fig. 5f). This characteristic input timing-dependence indicates that the plasticity mechanism enhanced by 4-AP or somatic depolarization is indeed related to ITDP, rather than some other form of long-term plasticity, such as LTP or spike timing dependent plasticity (STDP).

If dendritic spikes are critical for the induction of ITDP, we predicted that it might be possible to induce ITDP with fewer than 90 paired stimuli when dendritic spiking is enhanced. We therefore examined the effect of varying the number of paired stimuli on the magnitude of ITDP with the soma held at -70 mV in the absence and presence of 4-AP or with the soma held at -55 mV (no 4-AP). At a holding potential of -70 mV (in the absence of 4-AP), 80 stimulus pairs were required to produce a significant potentiation of the PSP measured 20 min after the induction protocol (p = 0.011 with Mann-Whitney test after Kruskal Walls test), with 90 stimulus pairs required to double the size of the PSP (an increase of 197.2 ± 19.6 %, n = 7, Fig. 6a,b). However, in the presence of 4-AP only 50 stimulus pairs were required to produce a significant potentiation (p = 0.039 with Mann-Whitney test after Kruskal-Wallis test), with 60 stimuli sufficient to cause a doubling of the PSP (PSP increase to 207.5 ± 27.3 % of its initial level, n = 5, Fig. 6a,c). Fifty stimulus pairs were also sufficient to cause a significant increase in the PSP in the presence of the Kv4.2/3 channel blocker phrixotoxin (the SC response increased to 200.4 ± 17.8 % after 20 min, p = 0.048, n = 4, Supplementary Fig. 6). The effect of somatic membrane depolarization to -55 mV was even more pronounced, with a significant potentiation being produced after only 20 stimuli (p = 0.031 with Mann-Whitney test after Kruskal-Wallis test), and 40 stimuli sufficient to more than double the PSP (223.8 ± 22.2 %, n = 5, Fig. 6a,d).

**Figure 6:**
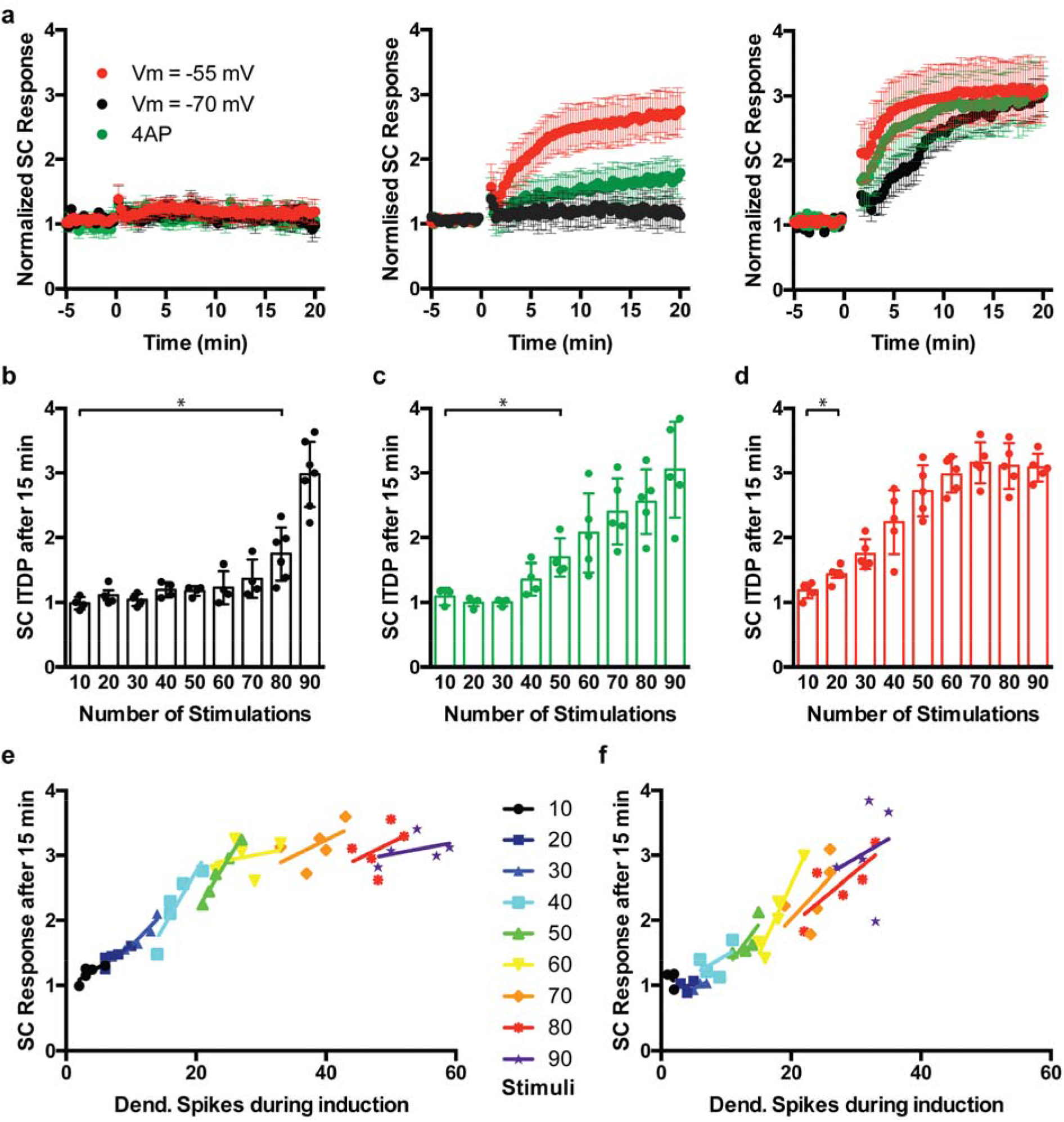
Somatic membrane depolarization or block of A-type K-channels enhances the efficiency of ITDP induction. **a** Change of the somatic PSP response to SC stimulation after ITDP induction, if the number of induction stimulus pairs is reduced to 10 (left), 50 (middle) or is kept at the usual 90 stimuli (right), colors indicate the condition of the cell during induction (black: Soma was held at -70 mV, green: 4-AP was present in the bath during induction, red: Soma was held at -55 mV during induction stimulation). **b** Average change in the somatic response to SC stimulation 20 minutes after ITDP induction, if the number of induction stimulus pairs is incrementally decreased from 90 pairs to 10 in steps of 10 and soma was held at -70 mV during induction. **c** Same as **b** but in the case that 4-AP was bath applied during ITDP induction. **d** Same as **b** but in the case that the soma was held at -55 mV during ITDP induction. **e** Correlation between number dendritic spikes during a given ITDP induction protocol and the increase in the SC response after 15 minutes when the soma is depolarized to -55 mV. **f** Same as **e)** but when the soma is held at -70 mV and 4-AP is present in the bath. For e and f, each point is an individual cell.

We observed a strong correlation between the number of dendritic spikes during the induction period and ITDP expression, especially over the range of 15-30 dendritic spikes. This was observed either when the soma was depolarized to -55 mV during ITDP induction (nonparametric Spearman correlation factor r = 0.66, p = 0.009 in this range, n = 15 cells, Fig. 6e) or when 4-AP was present in the bath (r = 0.78, p = 0.002, n = 14 cells, Fig. 6f). This correlation supports the view that dendritic spike firing during the paired synaptic stimulation plays an important role in the induction of ITDP. For technical reasons, most of the ITDP experiments were performed with only somatic recordings. As a result, we could not evaluate the correlation under control conditions, as dendritic spikes were only detectable in somatic recordings under conditions that favored spike propagation (see Fig. 4b).

Given the correlation between the number of dendritic spikes and ITDP, the effect of somatic depolarization to enhance the induction of ITDP is likely to result, at least in part, from the increased firing of dendritic spikes. Might the enhancement in dendritic spike propagation to the soma also contribute to the enhancement in induction of ITDP? To address this question we compared the relation between the onset of ITDP with the soma at either -70 mV or -50 mV as a function of the number of dendritic spikes observed during the induction protocol. We found that the onset of ITDP elicited by the same number of dendritic spikes was significantly greater when the membrane was depolarized. A linear fit to the ITDP versus spike number relation yielded y intercept values of -0.20 ± 0.43 at -70 mV and 0.35 ± 0.50 at -55 mV (p = 0.0005 with linear regression analysis; n = 7 (−70 mV) and 5 (−55 mV); Supplementary Fig. 5f). This result suggests that depolarization can enhance ITDP onset independently of enhancing dendritic spike firing, consistent with a role for DEDSP enhanced dendritic spike propagation. This change in elevation was not visible in the SC response 20 minutes after ITDP induction (data not shown), which is consistent with our earlier finding that after 20 minutes, the increase in the SC response reaches a plateau level (see Fig. 5).

As the dendritic spikes can trigger action potential output with the membrane held at -55 mV but not with the membrane at -70 mV, we examined whether the enhanced induction of ITDP from the more positive somatic potential was a result of the enhanced firing of somatic spikes. We locally applied TTX to the soma during paired stimulation to block the firing of somatic action potentials without altering dendritic spikes. Under these conditions, somatic depolarization was still able to enhance the induction of ITDP, which displayed a characteristic rapid onset and ability to be induced with only 50 paired stimuli (Supplementary Fig. 5). Similarly, even in the presence of TTX, there was a significant increase in the elevation of the correlation between the SC response, 2 minutes after ITDP induction and the number of dendritic spikes during induction (y intercept 0.34 ± 0.15, p = 0.0058 relative to -70 mV with linear regression analysis, n = 4, (Supplementary Fig. 5f)). These results clearly indicate that an increase in the efficacy of dendritic spike propagation and/or generation is sufficient to increase the efficacy of ITDP induction. However, as the rate of development of ITDP with the membrane at -55 mV is somewhat greater in the absence of TTX than in the presence of TTX, increased somatic spike firing is also likely to be important.

### Dendritic spikes are necessary and sufficient to induce ITDP in conjunction with SC stimulation

To probe more directly the importance of dendritic spikes, we determined whether they were sufficient and/or necessary to induce ITDP. We carried out dual patch clamp recordings at the soma and distal apical dendrite of CA1 PNs as before. However, instead of the normal ITDP induction protocol, we omitted the PP stimulus and administered 90 SC stimuli at a frequency of 1 Hz, either alone or paired with a dendritic spike elicited by direct dendritic current injection. The 90 SC stimuli alone caused no significant increase in the PSP in the following 20 minutes (113.5 ± 5.7 %, p = 0.65, n = 8, Fig. 7a-d). Next, we examined the effect, in the same cell, of pairing each of the 90 SC stimuli with the injection of a depolarizing 2-ms, 400-pA current step into the dendrite 3-5 ms after the SC stimulus. We chose this delay as it was within the time window in which we observed the onset of the dendritic spike during paired PP-SC stimulation at the -20 ms interval (see Fig. 4b upper traces) and roughly coincided with the peak of the SC PSP measured in the dendrite. The pairing of SC synaptic stimulation and distal dendritic current injection evoked a dendritic spike with a high probability (89.6%; n = 450 stimuli in 5 cells). Importantly, the pairing also induced a large ITDP-like enhancement in the SC PSP measured in the distal dendrite, which progressively increased in amplitude within the following 20 minutes to 194.8 ± 18.0 % of its value before pairing (from 6.23 ± 0.76 mV before pairing to 11.21 ± 0.84 mV after pairing, p = 0.008, n = 8, Fig. 7a-d).

**Figure 7:**
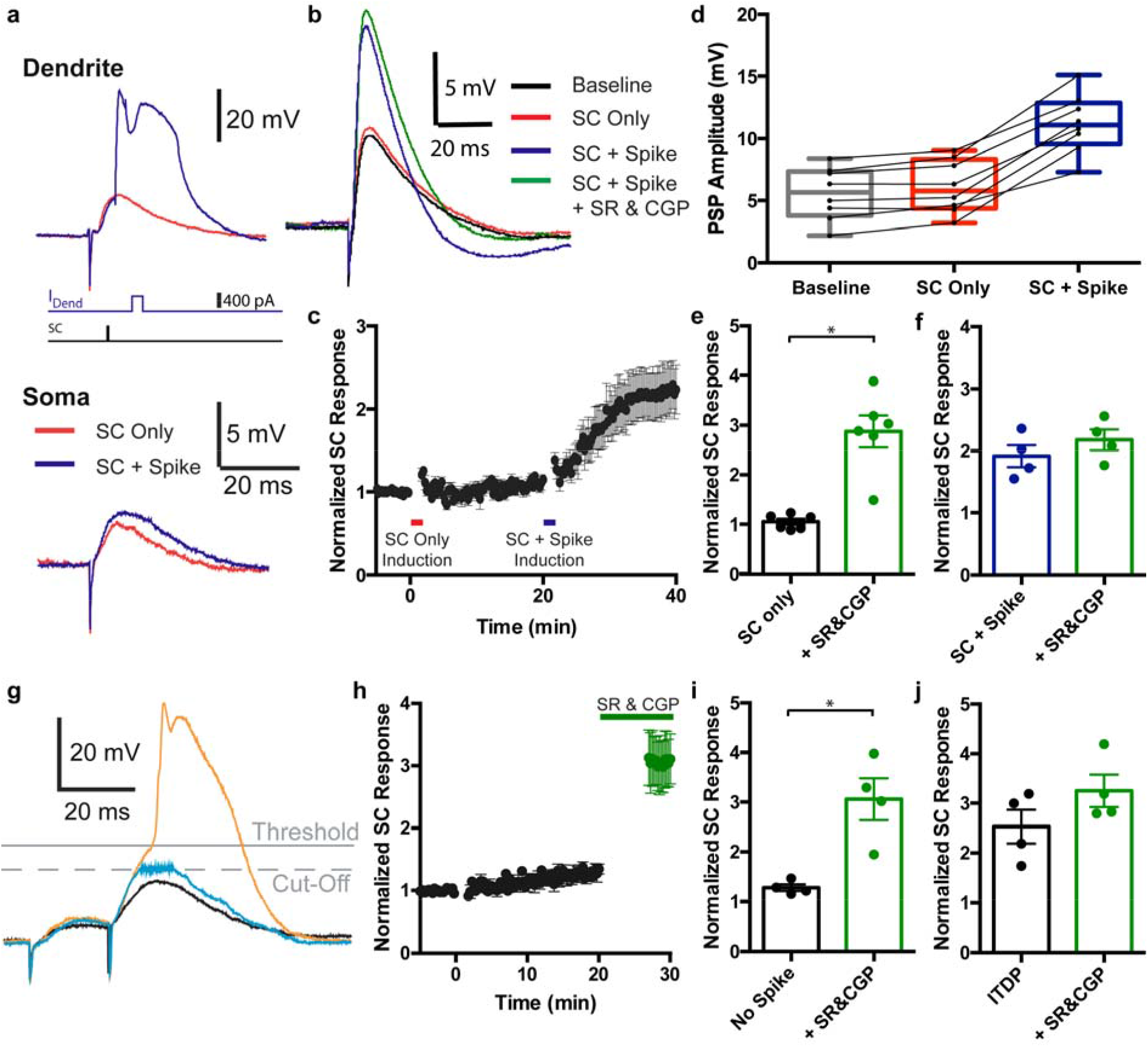
Dendritic spikes are necessary for ITDP induction. **a** Example traces of a modified ITDP induction protocol, where an SC stimulus is either combined with dendritic current injection (blue) or not (red), top: dendritic patch clamp recording, bottom: somatic recording. **b** The SC PSP of the same cell before any stimulation (black), 20 min after 90 single SC stimuli were administered at 1 Hz (red), 20 min after 90 SC stimuli were combined with dendritic current injection to elicit dendritic spikes at 1 Hz (blue) and after subsequent bath application of 2 μM SR95531 and 1 μM CGP55845 (green). **c** Time course of the change in the SC PSP over the course of the experiment, the bars indicate the time of the 1Hz SC stimulation first without (red) and then with (blue) dendritic current injection. **d** Summary of the SC PSP amplitude before and after the two stimulation modes described above, dots indicate single data points (n = 8). **e** Average SC PSP amplitude 20 min after SC stimulation only, before (black) and after (green) application of SR95531 and CGP55845 (n = 6). **f** Average SC PSP amplitude 20 min after combined SC stimulation and dendritic current injection before (blue) and after (green) bath application of SR95531 and CGP55845 in a subset of cells (n = 4). **g** Example traces of the dendritic voltage signal during ITDP induction: black: A PSP that does not reach dendritic spike threshold, orange: A PSP that does reach dendritic spike threshold as the custom script, used to block dendritic spikes electrically is disabled, light blue: A PSP that is prevented from reaching the threshold for a dendritic spike as negative current is automatically injected into the cell as soon as it reaches a voltage value, which is set below the dendritic spike threshold. **h** Time course of the change in SC PSP response during the time frame of ITDP expression when dendritic spikes are electrically blocked during ITDP induction, the green data points indicate measurements after bath application of SR95531 and CGP55845l. **i** Average change in SC PSP amplitude right before (black) and after (green) bath application of SR95531 and CGP55845 if dendritic spikes are electrically prevented during the induction protocol. **j** Same as **i** if dendritic spikes are allowed to happen during ITDP induction.

Does the potentiation induced by paired SC stimulation and dendritic spiking reflect the same plasticity mechanism as ITDP induced by paired synaptic stimulation? ITDP largely results from the long-term-depression of the component of feedforward inhibition evoked by Schaffer collateral stimulation (iLTD) that is mediated by cholecystokinin-positive (CCK^+^) interneurons (accounting for about 75% of the potentiation of the PSP), with a smaller component caused by LTP of the EPSP (eLTP; accounting for about 25% of the potentiation)^15^. To determine whether pairing of SC stimulation with dendritic spiking also suppressed inhibition, we assessed the extent of inhibition by applying blockers of GABA_A_ (2 μM SR95531) and GABA_B_ (1 μM CGP55845) receptors on the PSP amplitude before and after induction of plasticity. Normally, the application of the GABA receptor antagonists produced a nearly three-fold increase in the peak depolarization during the SC PSP because of blockade of the rapid feedforward IPSP (peak SC PSP increased to 282.6 ± 40.9 % of initial value, p = 0.031, n = 6, Fig. 7e). However, 20 min following the induction of ITDP with paired synaptic stimulation, the blockers increased the PSP to only 133.0 ± 14.2 % of its value relative to control (p = 0.125, n = 4, Fig. 7j), indicating a pronounced decrease in feedforward inhibition. We observed a similar loss of sensitivity to GABA receptor blockade 20 min after pairing SC stimulation with dendritic spikes, with the blockers increasing the PSP to only 114.3 ± 2.5% of control (PSP increased from 10.78 ± 1.29 mV to 12.35 ± 1.60 mV, p = 0.13, n = 4, Fig. 7f).

To further confirm that the plasticity induced by pairing SC stimulation with dendritic spike firing was related to ITDP, we applied the induction protocol with inhibition continuously blocked (using SR95531 and CGP55845). This largely eliminated the enhancement of the PSP, with only a small residual potentiation (127.5 ± 6.8 %, n = 4), which likely reflects the small component of eLTP recruited by ITDP induction (Supplementary Fig. 7a). These results thus support the view that the plasticity induced by pairing SC stimulation with dendritic spiking largely resulted from ITDP. In contrast, if the pairing protocol induced conventional Hebbian LTP or STDP, it should have been greatly enhanced with inhibition blocked (because of increased postsynaptic depolarization).

As a third approach to determine the nature of the plasticity mechanism, we examined the effects of bath application of the endocannabinoid CB1 receptor blocker AM251 (2 μM) during the ITDP induction protocol. Because the iLTD component of ITDP results from the activation of CB1 receptors on presynaptic terminals of CCK-expressing interneurons, AM251 blocks the majority of ITDP induced by paired synaptic stimulation^15^. Consistent with these results, application of AM251 also greatly reduced the synaptic potentiation produced by pairing dendritic spikes with SC stimulation, (144.3 ± 11.9 %, n = 5, which is significantly less than the potentiation in the absence of the blocker seen above; p = 0.0062), with the residual potentiation likely resulting from eLTP (Supplementary Fig. 7b).

We thus conclude that dendritic spikes in conjunction with SC stimulation are sufficient to induce ITDP. But are dendritic spikes necessary? Can ITDP be induced during paired distal and proximal synaptic stimulation if dendritic spikes are prevented? Addressing this question is complicated by the fact that pharmacological blockade of active dendritic events would also compromise synaptic transmission, which is essential for ITDP induction.

We therefore suppressed the generation of dendritic spikes electrically by using an online computer-controlled program to inject negative current into the dendrite to prevent dendritic spiking whenever the membrane depolarized to the dendritic spike threshold. This prevented dendritic spikes from occurring without interfering with summation of subthreshold distal and proximal PSPs. Prevention of dendritic spikes almost fully suppressed ITDP evoked by the standard paired synaptic stimulation protocol. Twenty minutes after delivering the induction protocol with dendritic spikes suppressed, the SC-evoked PSP was increased to only 126.5 ±7 % of its initial value, compared to the 274.3 ± 34.2 % increase under normal condition (p < 0.05 with Mann-Whitney test, n = 4; Fig. 7g-i, compare with Supplementary Fig. 4).

To assess the extent of inhibition when the ITDP induction protocol was delivered with dendritic spiking suppressed, we applied GABA_A_ and GABA_B_ receptor antagonists (SR95531 and CGP55845) 20 min after paired stimulation. After pairing, the blockers still caused a large increase in the net SC PSP amplitude (to 310.8 ± 40.0 %, p < 0.0001, n = 4, Fig. 7g,h,i), similar to their effect under baseline conditions. Thus, the prevention of dendritic spiking suppressed the induction of iLTD.

In contrast, when we allowed dendritic spikes to occur normally during the ITDP induction protocol, we observed an increase of the SC response to 252.9 ± 34.1 % of baseline (p = 0.045, n = 4), and subsequent application of SR95531 and CGP55845 caused only a slight, statistically insignificant increase in PSP amplitude (325.1 ± 32.6 % from baseline but only 133.0 ± 14.2 % from the post ITDP SC response, p = 0.125, n = 4, Fig. 7j), consistent with previous results showing that ITDP largely suppresses feedforward inhibition ^15^.

Together, these results indicate that dendritic spikes are both necessary and, in conjunction with SC stimulation, sufficient to induce ITDP.

## Discussion

Our results show that the propagation efficacy of dendritic spikes generated in distal CA1 dendrites is powerfully controlled by the somatic membrane potential. As a result of the effect of somatic depolarization to enhance dendritic spike propagation, the level of membrane polarization at the soma and proximal dendrite can regulate whether distal synaptic input that is strong enough to trigger a local dendritic spike will remain confined to the distal dendrite or will have a more powerful impact on signal integration and AP output at the soma. In this way DEDSP provides a powerful means by which synaptic input to one dendritic compartment can influence the efficacy of input to a distinct and spatially remote compartment.

Our results implicate dendritic A-type K^+^ channels as an important mechanism contributing to DEDSP. In ex vivo slices, neurons are relatively isolated from their surrounding network and synaptic input is sparse, so the resting membrane potential of CA1 pyramidal cells is stable and fairly negative, around -70 mV (an average of -68.9 ± 0.8 mV in these experiments, see Supplementary Fig. 1a). At this low membrane potential, A-type K^+^ channels will be in a fully available state with little resting inactivation^28^. As a result the channels will be able to conduct a large repolarizing K^+^ current in response to a subsequent depolarization. Thus, as a dendritic spike propagates actively from the distal dendrites towards the soma, these channels will rapidly activate and reduce the efficacy of spike propagation. However, if the cell is maintained in a slightly depolarized state prior to dendritic spike firing, a significant fraction of A-type channels in the proximal dendrites will be inactivated, decreasing the amount of outward K^+^ current activated by a dendritic spike and, thereby, permitting efficient active dendritic spike propagation. Our experimental results corroborate previous experimental results on the effect of membrane voltage to enhance the propagation of dendritic Na^+^ spikes along the apical dendrites of rat CA1 pyramidal neurons^22^. In addition, we find that resting potential powerfully regulates the calcium-dependent long-lasting dendritic spike, as the effect of somatic membrane potential on propagation remains in the presence of TTX (see Fig. 1 and 2). The results in this study are also consistent with experimental results showing that A-type K^+^ channels limit back propagation of action potentials into the dendrite^28, 38^.

Interestingly, the propagation of passive signals, such as single PSPs that did not reach the threshold for dendritic spiking, was not affected significantly by A-type K^+^ channels in the apical dendritic trunk. This may be due to the low amplitude of these signals, which may not cause sufficient A-type K^+^ channel activation to affect PSP amplitude attenuation at the soma.

A recent study by Hsu et al. (2018) has shown that the excitatory somatic response to a brief high frequency train of weak subthreshold distal EPSPs that fail to elicit a dendritic spike can also be amplified by somatic depolarization. However, here the mechanism does not involve enhanced dendritic propagation but rather results from a local boosting at the soma as a result of activation of a perisomatic persistent voltage-gated Na^+^ current^21^. Interestingly, Hsu et al. observed little effect of persistent Na^+^ current on single subthreshold EPSPs or actively propagating dendritic plateau potentials, similar to our results. Together, the report of Hsu et al. and our findings indicate that there are distinct activity-dependent mechanisms for enhancing the influence of distal PP inputs on CA1 pyramidal neuron output depending on the dynamics and strength of that input.

In addition to enhancing forward propagation of distally-generated dendritic spikes, we find that when the soma is in a more depolarized state, dendritic excitability in response to back-propagating action potentials is increased (Bock and Siegelbaum, unpublished). This effect is consistent with the known role of A-type K^+^ channels to limit backpropagating action potential (bAP) propagation ^28^. This would suggest that somatic membrane potential not only determines the impact of dendritic spikes on the soma but also regulates the efficacy of their initiation in response to somatic APs.

Our findings of DEDSP in acute hippocampal slices may help explain the efficient triggering of complex spikes when an animal occupies a pyramidal neuron’s place field. In vivo, CA1 pyramidal neurons receive a constant barrage of background synaptic inputs to their somato-dendritic compartment, leading to a more depolarized resting potential (around -60 mV) compared to the resting potential in acute slices (−70 mV) ^4, 19, 20, 39, 40, 41^. Moreover, in vivo recordings have shown that spike firing associated with place fields is typically preceded by a depolarizing ramp potential^39^, which may represent a combination of subthreshold EPSPs arising from inputs to basal or proximal apical dendrites and dendritic spikes that poorly propagate to the soma. As the resting potential depolarizes during the ramp, A-type K^+^ channel inactivation will increase progressively. Thus, place field membrane potential dynamics are ideally suited to enhance dendritic spike propagation and the firing of complex spikes.

DEDSP also provides a mechanism to explain recent in vivo findings that somatic depolarization through direct current injection can further enhance the frequency of complex spike firing ^19^ and lead to the appearance of de novo place field activity in a formerly silent CA1 neuron ^4, 20^. As we found that depolarization does not enhance propagation of subthreshold EPSPs, the ability of somatic depolarization to reveal the presence of subthreshold place field events suggests that these events likely result from distal dendritic spikes that are unpaired with proximal synaptic events. The coincidence of distal dendritic spikes with large somatic depolarization, either through direct somatic current injection or strong proximal synaptic input, triggers the firing of a long-lasting dendritic plateau potential ^4^, which results in a stable place field activity that outlasts the somatic depolarization, presumably through the induction of some form of synaptic plasticity.

Our results further identify the importance of dendritic spikes for the induction of one form of synaptic plasticity, ITDP. Thus, we both confirm that the ITDP induction protocol is able to trigger the firing of dendritic spikes (Basu et al.,2016) and now provide direct evidence that dendritic spikes are necessary, and, when paired with weak SC stimulation, sufficient to induce ITDP. Our study further reveals a marked influence of somatic membrane potential on the efficacy of ITDP induction. Thus, depolarization of the soma from its normal resting potential of -70 mV observed in acute slices to -55 mV, a value attained in vivo, decreases the number of synaptic pairing events required to produce a significant potentiation by a factor of 4 (from 80 to 20). This likely overestimates the minimum number of pairing events required to induce ITDP because the somatic depolarization achieved by current injection through a patch electrode will decrement in the dendrite as a function of distance from the soma. A more uniform depolarization of the entire somato-dendritic compartment, as achieved under physiological conditions, may enable ITDP induction with even fewer paired events.

Somatic depolarization is likely to enhance ITDP through several mechanisms. First, we find that somatic depolarization enhances the ability of the dendritic spikes to trigger a somatic action potential, which also increases the firing of bAPs. As bAPs can broaden the dendritic spike^42^, they will enhance Ca^2+^ influx, which could help promote ITDP. However, since we find that somatic depolarization significantly enhances ITDP even when somatic action potentials and bAPs are blocked by local somatic application of TTX, other processes must also contribute. One likely additional mechanism is through the effect of somatic depolarization to increase the total number of dendritic spikes elicited by the synaptic pairing protocol, as we find that ITDP depends on the number of dendritic spikes. Finally, somatic depolarization is also likely to enhance ITDP by enhancing dendritic spike propagation, irrespective of the increase in number of dendritic spikes. This conclusion is supported by our finding that when the resting membrane was depolarized a given number of dendritic spikes led to an increased onset of ITDP compared to when the membrane was at its normal negative resting potential. All the above factors likely combine to result in a greater depolarization at the proximal dendrites and soma to enhance the perisomatic Ca^2+^ influx. Based on previous results, the enhanced Ca^2+^ influx synergistically interacts with the activation of metabotropic glutamate receptors to trigger endocannabinoid release, which leads to the induction of ITDP^9, 15, 16^.

The effect of somatic depolarization to increase the probability of distal dendritic spike firing during the ITDP induction protocol is surprising, given that somatic depolarization produces only a small depolarization in the distal dendrite. Moreover, we find that this small distal depolarization is insufficient to alter the threshold of dendritic spiking in response to direct depolarizing current injection in the distal dendrite. One clue as to the potential mechanism for enhanced firing of dendritic spikes comes from our finding that the increase in the probability of dendritic spike firing during ITDP induction is not seen with the first few pairs of synaptic stimuli but only develops after several pairing events. This suggests that the pairing protocol gives rise to some form of activity-dependent intrinsic or synaptic plasticity process that enhances dendritic spiking, and that this plasticity mechanism is enhanced by somatic depolarization. Although further studies will be required to identify the basis of this plasticity, one simple hypothesis is that the plasticity itself is driven by dendritic spike firing, and that somatic depolarization enhances the plasticity by enhancing dendritic spike propagation to the soma.

Another potential mechanism by which somatic depolarization could, in principle, enhance dendritic spike propagation is through the enhancement of NMDA receptor opening and/or the firing of NMDA spikes. However, several lines of evidence suggest that enhanced NMDA receptor activation does not mediate the effect of somatic depolarization to enhance either dendritic spike propagation or ITDP induction. Thus, we found that DEDSP was observed even when a distal dendritic spike was directly triggered by dendritic current injection, and thus independent of NMDA receptor activation (see Fig. 1 and 2). Furthermore, we found that DEDSP occurred when only distal PP synapses were stimulated (Fig. 3). As somatic depolarization produced little change in resting potential in the distal dendrites, the sole site of NMDA receptor activation in these conditions, enhanced NMDA receptor activation is unlikely to account for enhanced dendritic spike propagation. Finally, a recent study investigating passive signal propagation in CA1 PN dendrites found that blocking NMDA receptors has only marginal effects on the somatic voltage-dependent amplification of single EPSPs or trains of EPSP ^21^.

Although we cannot directly test whether the effect of somatic depolarization to enhance ITDP results from enhanced NMDA receptor activation at the SC synapses, because NMDA receptors are required for the induction of ITDP^16^, we also think it unlikely for the following reasons. First, the depolarization in the proximal dendrite in response to a 15 mV somatic depolarization is small (<5-10 mV), and much less than the 70 to 100 mV depolarization typically required to induce classic NMDA receptor dependent plasticity^43^. Second, somatic depolarization paired with a SC-evoked EPSP alone (without a dendritic spike) failed to induce plasticity under the conditions of our experiment, suggesting minimal Ca^2+^ influx or plasticity absent a dendritic spike. Finally, our finding that closed loop blockade of dendritic spikes with subthreshold EPSPs left intact was sufficient to suppress ITDP suggests that Ca^2+^ influx via NMDA receptors alone is insufficient to induce plasticity.

In addition to ITDP, we speculate that other NMDA receptor dependent forms of plasticity that require coordinated activity in multiple subcellular compartments may also be enhanced as the soma is depolarized. Spike timing dependent plasticity (STDP) is one such example, where dendritic excitability in conjunction with bAPs promotes synaptic plasticity ^44, 45^. Given that both dendritic excitability and back-propagation of sodium spikes are modulated by membrane potential, a more depolarized cell may generate a more efficient STDP, potentially with a longer time window for plasticity to occur. Furthermore, the additional somatic depolarization that DEDSP provides may in turn cause additional somatic APs (as e.g. seen in figure 4), which have an impact on plasticity mechanisms as well.

As many factors contribute to determining the somatic membrane potential, our results have a number of additional implications. First, what was formerly perceived as subthreshold background activity in neurons in vivo can, by influencing resting potential, powerfully regulate a neuron’s spiking output and tune its sensitivity towards plasticity. This will serve to increase a neuron’s computational capability at the subthreshold level as a given input, even if it does not contribute to a spike directly, will be able to modulate neural responsiveness throughout a neuron’s complex dendritic arbor, even to events that arise in a different subcellular compartment. Such a mechanism, for example, would allow subthreshold input at a neuron’s basal dendrites to enhance spike propagation of distal inputs along the apical dendrite. Second, as neuromodulatory transmitters often act to regulate resting potential, DEDSP may also contribute to effects of arousal and attention on memory storage and sensory perception. Future in vivo studies will be of interest to examine how dendritic spike propagation is regulated during relevant behaviors and how direct manipulations of dendritic spike propagation efficacy may alter those behaviors.

## Materials and Methods

### Slice preparation

C57-BL6 Mice (7-12 weeks old) were anesthetised by inhalation of isoflurane (5%) for 7 minutes, subjected to cardiac perfusion of cold ASCF for 30 seconds before decapitation according to the procedures approved by the IACUC of Columbia University and the New York State Psychological Institute. The skull was opened and the brain removed and immediately transferred into ice-cold carbogenated artificial cerebrospinal fluid (ACSF; 22.5 mM glucose, 125 mM NaCl, 1 mM MgCl_2_, 2 mM CaCl_2_, 25 mM NaHCO_3_, 3 mM KCl, 1.25 mM NaH_2_PO_4_, pH=7.2). During the slicing procedure, the MgCl_2_ concentration was increased to 4 mM to reduce cell excitability. The hippocampus was dissected in both hemispheres. 400 µm thick horizontal hippocampal slices were cut using a vibrating tissue slicer (VTS 1200, Leica, Germany) and transferred to a chamber containing ACSF at 35°C, where they were incubated for 40 minutes. Thereafter, slices were held at room temperature (21°C) until transfer into the recording chamber.

### Electrophysiology

Slices were transferred from the incubation chamber into the recording chamber of an Olympus BX51WI microscope (Olympus, Japan), where they were either held in place by a 1mm grid of nylon strings on a platinum frame or – in the case of experiments that required dual extracellular stimulation – by pinning the slice to the bottom of the chamber with the two stimulation electrodes. Slices were continually perfused with ACSF at a temperature of 34 ± 1°C, maintained by a thermostat controlled flow-through heater (Warner Instruments, CT)

In the case of single somatic recordings, healthy somas of CA1 pyramidal neurons were identified visually under 80x (40×2) magnification and patched under visual guidance using borosilicate glass pipettes (i. d. 0.75 mm, o. d. 1.5 mm, Sutter Instruments, UK) with a tip resistance of 4-5.5 MΩ, connected to a Multiclamp 700B amplifier (Molecular Devices, CA) and filled with intracellular solution, containing (in mM): 135 K-gluconate, 5 KCl, 0.2 EGTA, 10 HEPES, 2 NaCl, 5 MgATP, 0.4 Na_2_GTP, 10 Na_2_-Phosphocreatin, adjusted to a pH of 7.2 with KOH. Recordings were only accepted if the series resistance after establishing a whole cell configuration did not exceed 22 MΩ and did not change by more than 20% of the initial value during the course of the experiment.

In the case of single dendritic patches, a thick walled borosilicate glass pipette (i.d. 0.75 mm, o.d. 1.6 mm) with a tip resistance of 9 – 13 MΩ was lowered slowly (<2 µm/sec) into the distal Stratum Radiatum of CA1. Dendrites were patched blindly by tracking changes in the tip resistance. Recordings were included into the analysis if the series resistance after establishing whole cell configuration did not exceed 45 MΩ and if it did not change by more than 20% of the initial value during the course of the experiment.

In the case of dual dendritic and somatic patch clamp recording, both methods described above were combined, starting with the dendritic blind patch. In this case, 5 µM Alexa 594 fluorescent dye (ThermoFisher Scientific, MA) was added to the intracellular solution of the dendritic patch pipette in order to visualize the soma of the same cell and target it for the somatic patch clamp.

In the case of localized extracellular synaptic stimulation, stimulator pipettes were pulled from borosilicate glass capillaries (i. d. 0.75 mm, o. d. 1.5 mm, Sutter Instruments, UK) with a tip resistance of 50-70 MΩ. These pipettes were filled with 1M NaCl solution, attached to electrode holders, connected to a separate grounding electrode in the bath and inserted into the slice in stratum radiatum (SR) (for SC stimulation) or SLM (for PP stimulation) at a distance of at least 180 µm from the patched cell and/or dendrite in order to avoid direct electrical stimulation of the patched cell. Stimulations were administered through battery powered stimulator boxes, connected to the stimulation and ground electrodes and triggered via TTL signals.

### Pharmacology

Drugs were prepared as stock solutions and stored until needed. They were then diluted to final concentration in ACSF and added to the bath. TTX (Sigma-Aldrich, MO) was stored as a stock solution of 1 mM in ACSF at -20°C and applied at 500 nM to the bath. Cd^2+^ (Sigma-Aldrich, MO) was stored as a stock solution of 100 mM CdCl_2_ in ACSF at 4°C and applied to the bath at 200 μM. 4-AP (Tocris, UK) was stored as a stock solution of 100 mM in ACSF at -20°C and applied to the bath at 5 mM. Phrixotoxin 2 (Alomone Labs, Israel) was stored at a stock concentration of 5 µM and bath applied at a concentration of 500 nM. SR95531 (Tocris, UK) was stored as a stock solution of 20 mM in ACSF at - 20°C and applied to the bath at a concentration of 2 μM. CGP55845 (Tocris, UK) was stored as a stock solution of 10 mM at -20°C and applied to the bath at 1 μM. AM251 (Tocris, UK) was stored in a stock solution of 20 mM in DSMO and added to the bath at a concentration of 2 μM.

### Data acquisition and analysis

Electrophysiological recordings were digitized, using a Digidata 1322A A/D interface (Molecular Devices, CA), at a sampling rate of 20 kHz (low pass filtered at 10 kHz) and recorded pClamp 10 software (Molecular Devices, CA). The amplifier setting of the Multiclamp 700B were controlled through Multiclamp Commander (Molecular Devices, CA). In one experiment (Fig. 7g-i) a custom designed script, built on the basis of Ephus software ^46^ was used to perform online analysis and insert a negative current into the cell, based on the membrane voltage (see Fig. 7G).

Data was analyzed using Axograph X software (Axograph Scientific, Australia), MATLAB (Mathworks, MA) as well as Microsoft Excel (Microsoft Corp., WA). Prism 4/5 (Graphpad, CA) was used for statistics and preparation of graphs. For paired data Wilcoxon’s non-parametric matched pairs test or a paired t-test (if a Gaussian distribution could be assumed) was used to test statistical significance unless otherwise noted. For multiple data sets Dunn’s Multiple Comparison test was performed to determine statistical significance unless otherwise noted. Results are presented as average values ± standard error (SE). Data was visualized in Acrobat Illustrator (Adobe, CA). Example traces represent raw data recordings with electrical artifacts of extracellular stimulation downscaled to 3-15%.

## Acknowledgements

We would like to thank the members of the Siegelbaum lab for helpful discussions. This work was funded by the National Institute of Health.

## Author Contributions

T.B. and S.A.S. conceptualized and designed the experiments, T.B. conducted the experiments and data analysis, T.B. and S.A.S. wrote the manuscript.

## Data availability

The datasets generated and analyzed during the current study are available at: https://drive.google.com/drive/folders/1ANHQhYiUdb6jvS-NzBVnqwINx5ry5YfA?usp=sharing

