## Supplemental Figures for "Somatic Depolarization Enhances Hippocampal CA1 Dendritic Spike Propagation and Distal Input Driven Synaptic Plasticity"

### Supplemental Material:

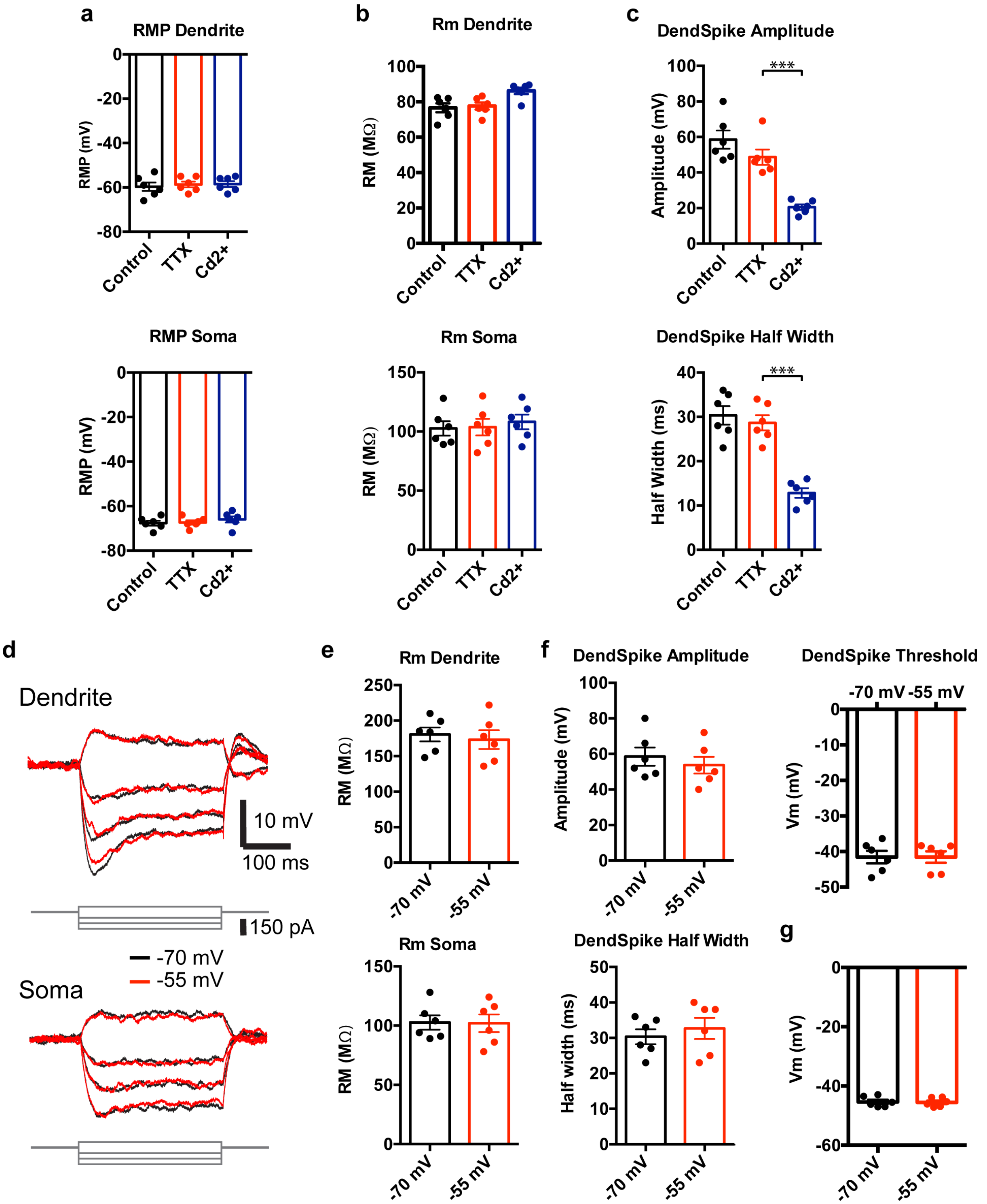

**Supplementary Figure 1:** Passive membrane properties and dendritic spike properties do not significantly change in response to TTX or depolarization. **a** Resting membrane potential at the soma (bottom) and the dendrite (top), 10 minutes after break-in (black) and after application of 500 nM TTX (red) and 200 µM Cd (blue). **b** Membrane resistance at the soma (bottom) and the dendrite (top), 10 minutes after break-in (black) and after application of TTX (red) and cadmium (blue). **c** Dendritic spike properties (top: Amplitude, bottom: half width), 10 minutes after break-in (black) and after application of TTX (red) and cadmium (blue). **d** Example traces of measurements used for the calculation of passive membrane properties at the soma (bottom) and dendrite (top). The cell is either held at -70 mV (black) or -55 mV (red), baselines were adjusted for better comparison. **e** Membrane resistance at the soma (bottom) and the dendrite (top) with the cell held at -70 mV (black) or -55 mV (red). **f** Dendritic spike properties with the cell held at -70 mV (black) or -55 mV (red). Top left, spike amplitude; bottom left, spike half-width; top right, spike threshold. **g** Somatic AP threshold with the cell held at -70 mV (black) or -55 mV (red).

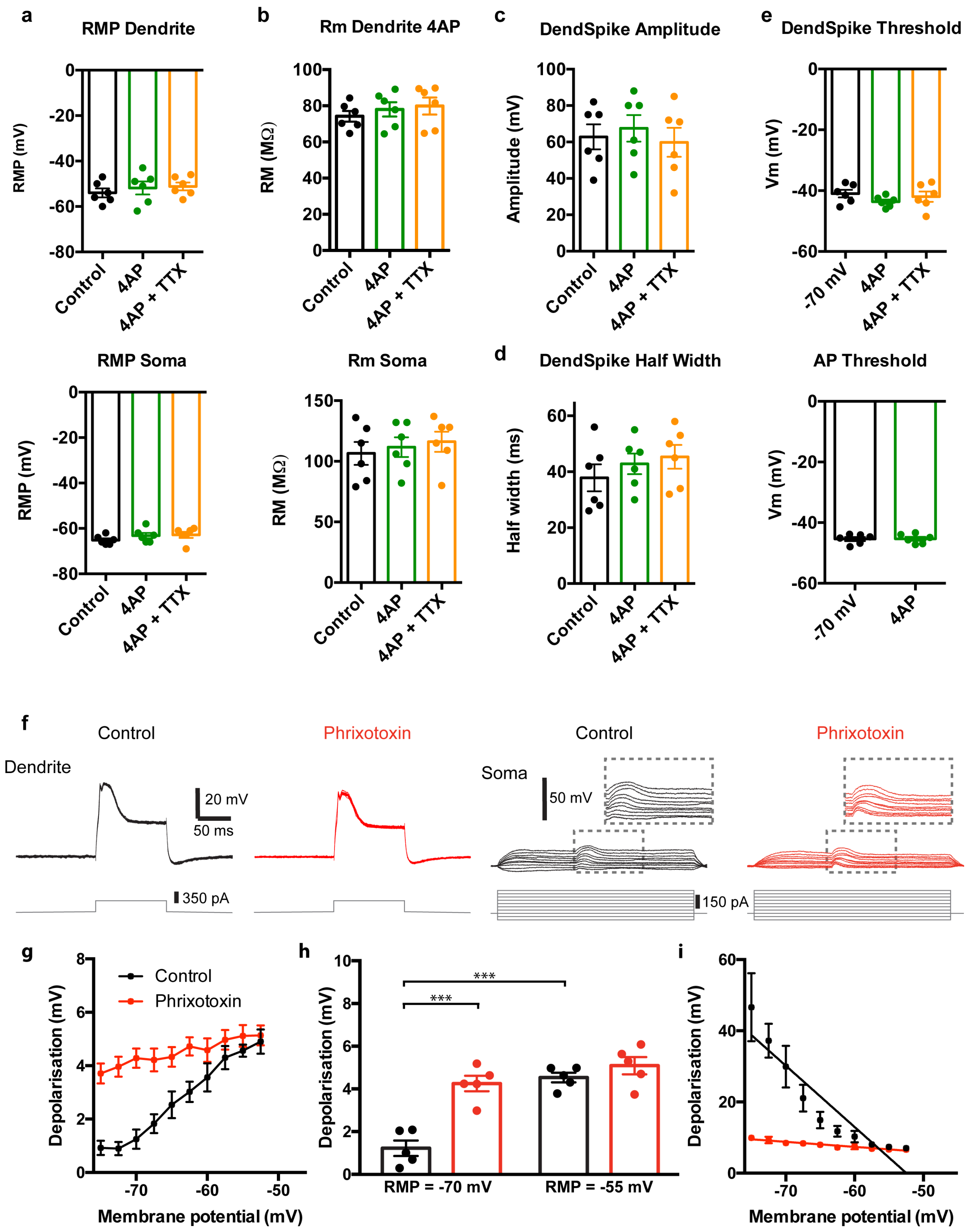


**Supplementary Figure 2:** Passive membrane properties and dendritic spike properties do not significantly change in presence of 4-AP or phrixotoxin. **a** Resting membrane potential at the soma (bottom) and the dendrite (top), 10 minutes after break-in (black) and after application of 5 mM

4-AP (green) and 500 nM TTX (orange). **b** Membrane resistance at the soma (bottom) and the dendrite (top), 10 minutes after break-in (black) and after application of 4-AP (green) and 500 nM TTX (orange). **c, d** Dendritic spike properties (top: Amplitude, bottom: half width), 10 minutes after break-in (black) and after application of 4-AP (green) and 500 nM TTX (orange). **e** Dendritic spike (top) and somatic AP (bottom) threshold 10 minutes after break-in (black) and after application of 4-AP (green) and 500 nM TTX (orange). **f** Dendritic (left) and somatic (right) recording of a dendritic spike, elicited by current injection into the dendrite at varying somatic membrane potentials before (black) and after adding 500 nM phrixotoxin (red) to the bath solution. Inserts show an expanded view of the somatic voltage responses of regions in dashed boxes. g Peak depolarization of the soma during the dendritic spike at varying somatic baseline potentials before (black) and after adding phrixotoxin (red) to the bath. **h** Comparison of peak somatic membrane depolarization with -70 mV and -55 mV baseline somatic membrane potentials before (black) and after adding phrixotoxin (red) to the bath. **i** Attenuation factor of the dendritic calcium spike, as it propagates to the soma, plotted against the somatic baseline potential before (black) and after phrixotoxin (red) to the bath.

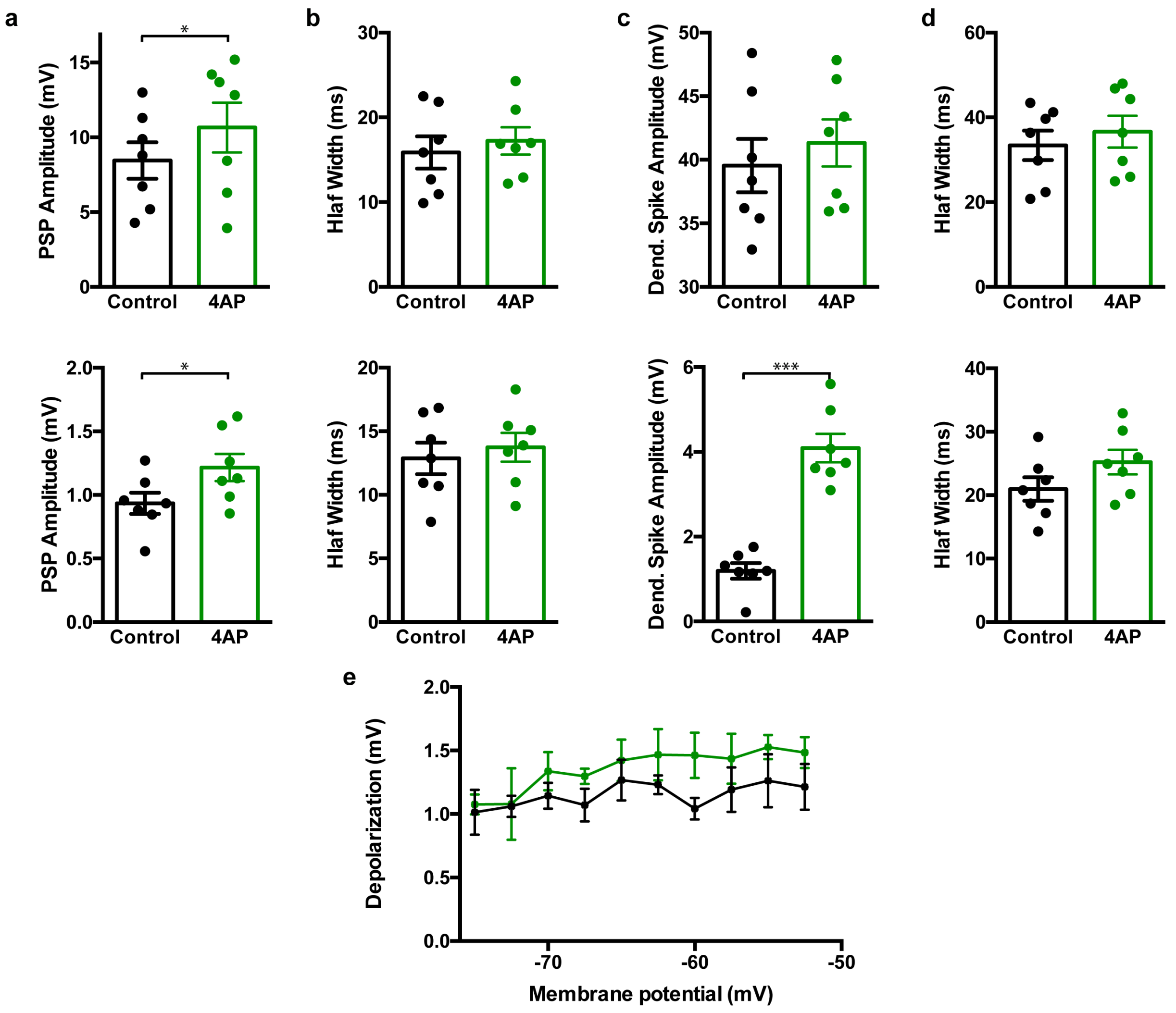


**Supplementary Figure 3:** Changes in amplitudes and duration of PSPs and dendritic spikes in the presence of 4-AP. **a** Amplitude of passively propagating PSPs, evoked by single PP stimulation at the dendrite (top) and the soma (bottom) before (black) and after(green) bath application of 5mM 4-AP. **b** Half width of passively propagating PSPs, evoked by single PP stimulation at the dendrite (top) and the soma (bottom) before (black) and after(green) bath application of 4-AP. **c** Amplitude of actively propagating dendritic spikes, evoked by high frequency PP stimulation at the dendrite (top) and the soma (bottom) before (black) and after(green) bath application of 4-AP. **d** Half Width of actively propagating dendritic spikes, evoked by high frequency PP stimulation at the dendrite (top) and the soma (bottom) before (black) and after(green) bath application of 4-AP. **e** Peak depolarization of the somatic membrane potential during a passively propagating PSP, evoked by direct dendritic current injection, mimicking a PSP waveform at varying somatic baseline potentials before (black) and after adding 4-AP (green).

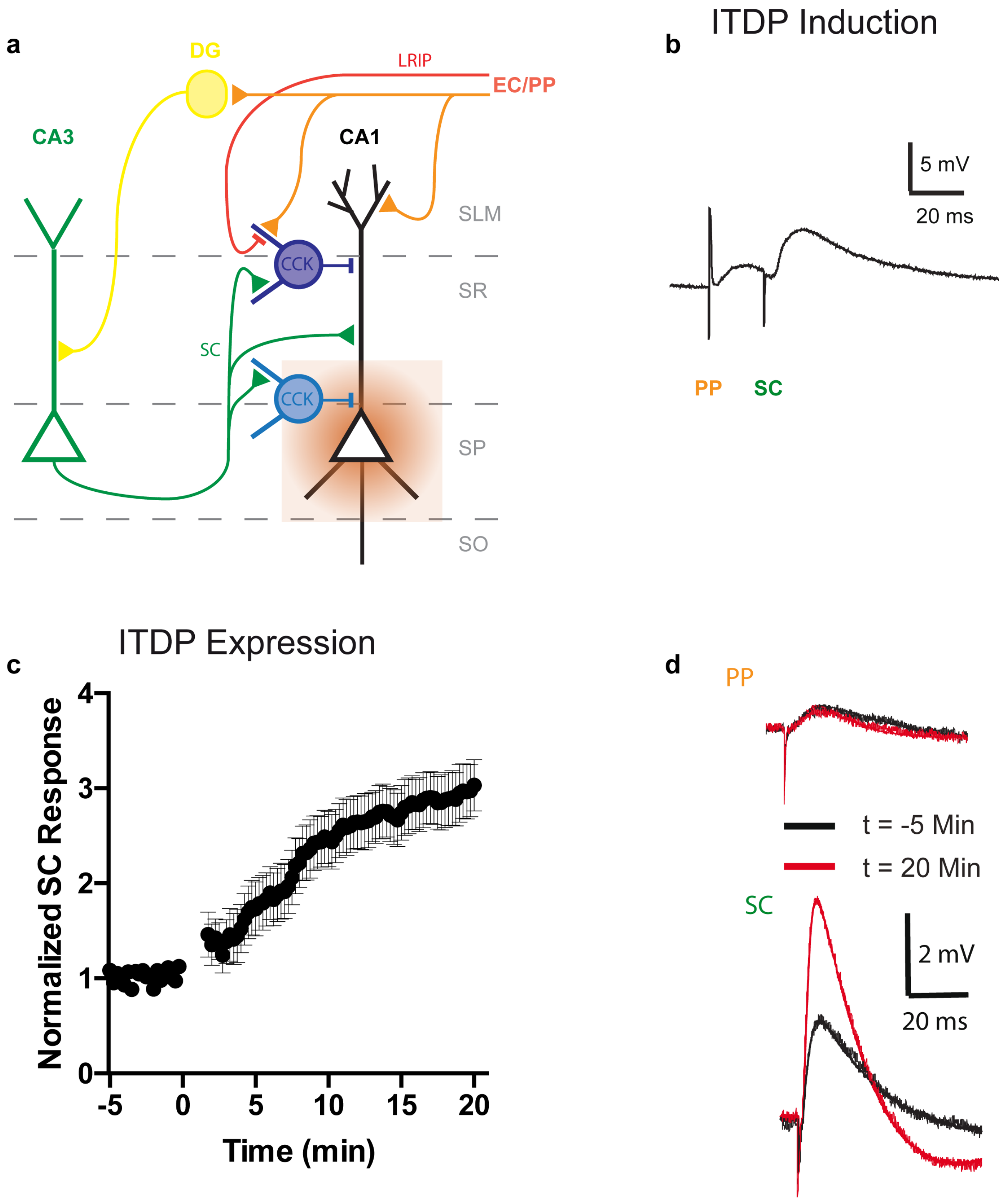


**Supplementary Figure 4:** ITDP Induction and expression. **a** Schematic of the ITDP circuit. Black: CA1 pyramidal neuron; green: CA3 pyramidal neuron, projecting to CA1 via Schaffer collaterals; orange: excitatory PP connections from EC neurons; red: long range inhibitory connections, also within the PP, coming from EC; yellow: Dendate Gyrus neurons; purple: dendrite targeting CCK interneurons in SR and SLM of CA1; blue: proximal and somatic targeting CCK interneurons, located in proximal SR and stratum pyramidale (SP); brown: endocannabinoids are released in the somatic and proximal compartment due to intracellular calcium influx in the soma of CA1 pyramidal neurons. **b** Example trace of a somatic patch clamp recording during an ITDP induction stimulus, the PP stimulus precedes the SC stimulus by 20 ms, this stimulus is repeated 90 times at 1 Hz. **c** Time course of ITDP expression, within 20 minutes after induction, the SC response increases by 197.2 ± 19.6 %, (n = 7). **d** Example traces of PP and SC stimulus responses 5 minutes (black) and 20 minutes (red) after ITDP induction.

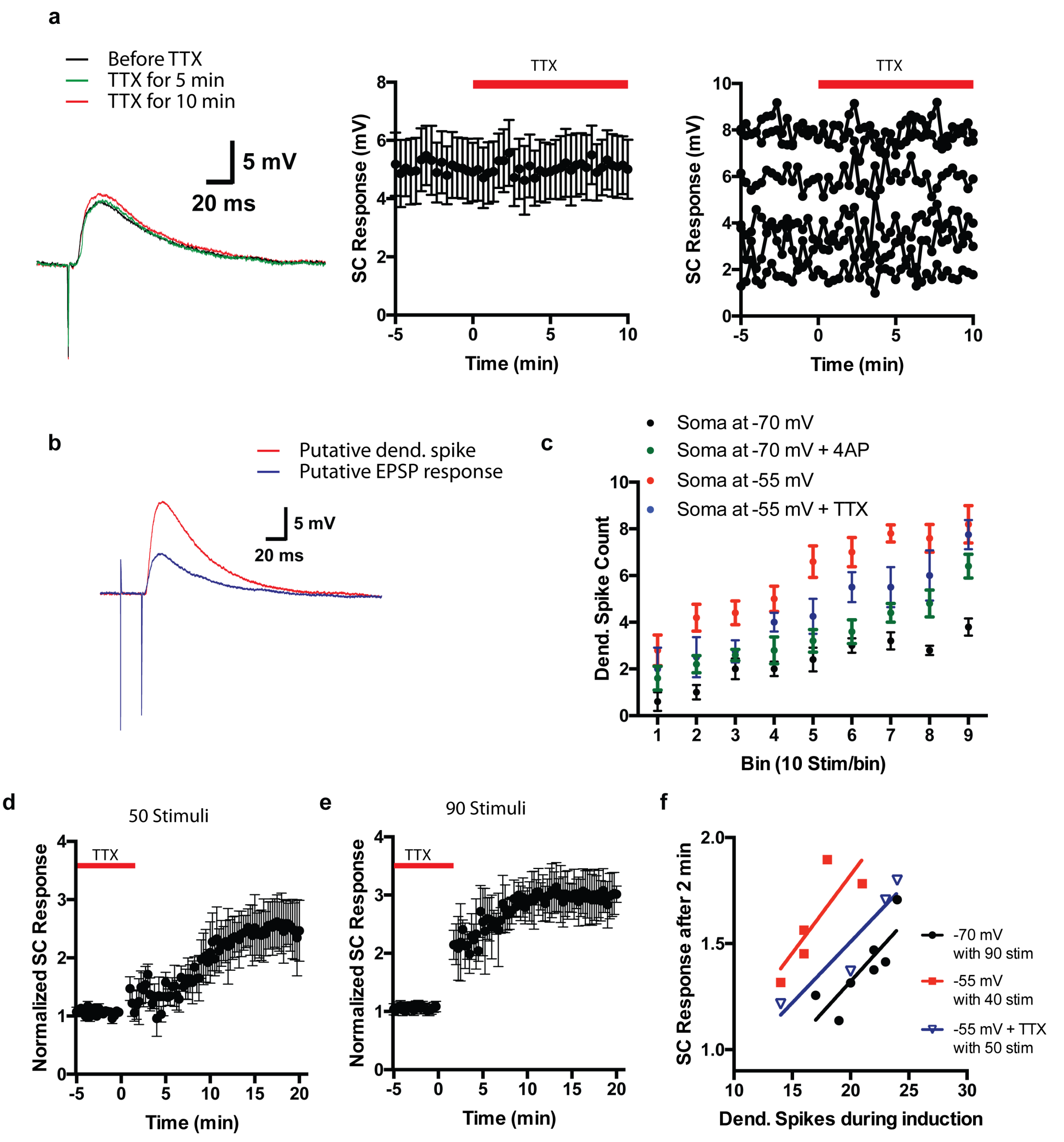


**Supplementary Figure 5:** Depolarization of the somatic membrane during ITDP induction in the presence of TTX. **a** Left: Example traces of a single SC stimulus to a CA1 pyramidal neuron before (black), after 5 min (green), and after 10 min (red) of TTX, applied to the soma through a local puffing pipette, Middle: The average response to SC stimulation during local somatic TTX application remains unchanged (n = 6), Right: Same as the middle graph but showing the responses of the individual cells. **b** Example traces of the somatic response to ITDP induction stimulation in those cases where a dendritic spike occurred (red) or not (blue) while the cell soma was depolarized to -55 mV and while TTX was applied to the soma (compare to figure 4b). **c** Average occurrence of dendritic spikes during the induction period, the 90 paired stimuli were separated into 9 bins of ten stimuli each and the number of dendritic spikes in each bin was counted (black: Soma was held at -70 mV, red: Soma was held at -55 mV during induction stimulation, green: 4-AP was present in the bath during induction, blue: TTX was applied to the soma and the soma was held at -55 mV during induction stimulation). **d** Change of the somatic PSP response to SC stimulation after 50 ITDP induction stimuli when TTX was applied to the soma and the soma was depolarized to -55 mV during induction (n = 4)**. e** Same as **d** but after 90 induction stimuli (n = 3). **f** Correlation between the total number of dendritic spikes during ITDP induction and the SC response 2 minutes after ITDP induction (normalized to PSP before induction) within a stimulus range that yielded a total of 13-27 dendritic spikes when the cell soma was held at -70 mV (black), or at -55 mV under control conditions (red) or during somatic application of TTX (blue).

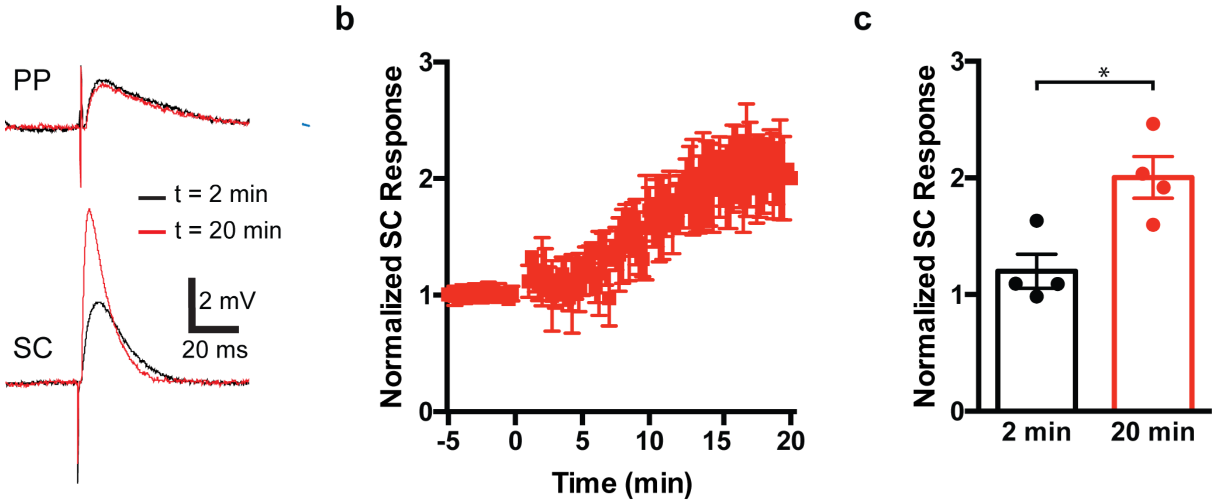


**Supplementary Figure 6:** ITDP after 50 stimuli in the presence of phrixotoxin. **a** Example traces of PSPs in response to perforant path (PP, top) and SC stimulation (bottom) 2 minutes (black) and 20 minutes (red) after 50 ITDP induction stimuli in the presence of 500 nM phrixotoxin in the bath solution. **b** Time course of ITDP expression, within 20 minutes after induction (n = 4 cells). **c** Average SC response 2 minutes (back) and 20 minutes (red) after ITDP induction (n = 4 cells).

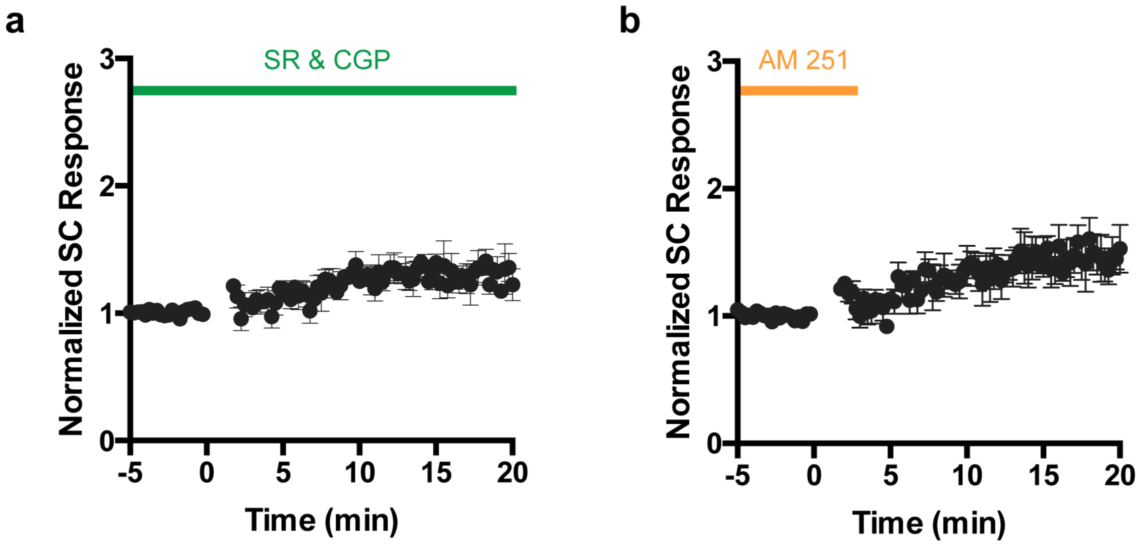


**Supplementary Figure 7:** Corroborating evidence that SC stimulation in conjunction with dendritic current injection to elicit dendritic spikes leads to ITDP, rather than LTP expression. **a** Time course of the change in SC PSP response after inducing ITDP with 90 SC stimuli at 1 Hz, each one combined with dendritic current injection to elicit dendritic spikes in the presence of 2 μM SR95531 and 1 μM CGP55845, the plasticity effect is occluded when interneurons are blocked as it depends on disinhibition, green bar indicates the time at which SR95531 and CGP55845 are present in the bath (n = 4). **b** Time course of the change in SC PSP response after inducing ITDP with 90 SC stimuli at 1 Hz, each one combined with dendritic current injection to elicit dendritic spikes after 2 μM of AM251 were present in the bath during the stimulation, the plasticity effect is blocked as the down regulation of interneurons that causes ITDP is mediated by presynaptic endocannabinoid receptor activation , orange bar indicates the time at which AM251 was present in the bath (n = 5).
